# Serum Lipidome as an Early Peripheral Indicator in Familial Alzheimer’s Disease

**DOI:** 10.1101/2025.11.25.690392

**Authors:** Gloria Patricia Cardona-Gómez, Iván Daniel Salomón-Cruz, Laura Alejandra Lozano-Trujillo, Nelson David Galvis-Garrido, Sara Camila Agudelo-Castrillon, Juan Pablo Barbosa-Carvajal, Mateo Hoyos Ríos, Lina M. Trujillo-Chacón, Eliana Henao, Geysson Javier Fernandez, Edison Osorio, Yakeel T. Quiroz, Eric Schon, Francisco Lopera, Andrés Villegas, Daniel Camilo Aguirre-Acevedo, Julián D. Arias-Londoño, David Aguillón, Estela Area-Gómez

## Abstract

Protein biomarkers in biofluids are highly sensitive indicators of prodromal cognitive impairment yet remain limited for primary prevention. Lipids, essential to brain structure and function, offer untapped prognostic value. Here, we identify a lipidomic signature in serum from asymptomatic PSEN1-E280A mutation carriers aged 6–40 years, that differentiate carriers from non-carriers with an AUC 80–90%. Similarly, to symptomatic carriers (≥41 years; 93%) and sporadic AD cases (85%), using high-resolution mass spectrometry. Latent profile analysis revealed lipid-based signatures of dementia risk and resilience, shaped by genotype, sex, and APOE isoform, and supported by SIMOA protein biomarkers. Age-dependent dysregulation in sphingolipid and glycolipid metabolism was validated by enzymatic activity (TLC), glial phenotyping (flow cytometry), and gene expression (snRNAseq) in postmortem brain. Ganglioside clearance deficits emerged by age 6–12, followed by proinflammatory shifts from age 13 and p-tau217 elevation by age 20, with greater burden in females and APOE4 carriers. APOE3Ch individuals showed differential salvage pathways of ceramides and gangliosides. These findings position early lipid pathway dysregulation as a biological contributor to Alzheimer’s pathogenesis and a potential therapeutic target for primary prevention.

## INTRODUCTION

Alzheimer’s disease (AD) is a progressive neurodegenerative disorder with a preclinical phase that begins decades before the symptoms appear. The conceptual shift from clinical to biological definitions of AD, articulated in the ATN framework, which refers to amyloidosis, tauopathy and neurodegeneration, has enabled researchers to characterize disease progression through measurable biomarkers in cerebrospinal fluid and blood, including Aβ₁₋₄₂, phosphorylated tau (p-tau), and neurofilament light chain (NfL)^1–4^. These markers delineate the molecular trajectory from silent pathology to cognitive impairment and now serve as essential tools for early detection and disease staging. However, their invasive collection, high cost, and limited scalability restrict their use for population-level screening and prevention strategies, particularly in low-resource settings.

Plasma-based biomarkers, such as p-tau217, have emerged as promising non-invasive alternatives with high specificity for amyloidosis and tauopathy^5–7^. However, they remain influenced by biological variability and temporal fluctuations relative to central pathology^7^, and they do not fully capture the metabolic processes that contribute to neuronal vulnerability. Therefore, identifying stable, peripheral biomarkers that reflect early pathophysiological alterations rather than later stages of protein aggregation, represents a critical gap in current Alzheimer’s disease research.

Among the early mechanisms under investigation, changes in lipid homeostasis have gained recognition as a central feature of neurodegeneration. Lipids account for nearly half of the brain’s dry weight and play critical roles in membrane structure, myelination, neurotransmission, and synaptic plasticity. Alterations in the metabolism of phospholipids, sphingolipids, ceramides, cholesterol, and polyunsaturated fatty acids (PUFA) contribute to amyloid aggregation, tau hyperphosphorylation, neuroinflammation, and synaptic dysfunction^8–11^. In this context, lipidomic profiling offers a powerful window into systemic biochemical changes, potentially revealing early signatures that precede irreversible neuronal loss.

Evidence from both central and peripheral lipidomic studies supports this perspective. Decreases in phosphatidylcholine (PC) and phosphatidylethanolamine (PE) species containing docosahexaenoic acid (DHA) have been observed in CSF and plasma, and impairment of lipid transport associated with the APOE ε4 genotype, which correlates with increased Aβ aggregation and earlier disease onset^12–15^. Importantly, the influence of APOE isoform background extends beyond ε4-associated risk, as protective variants such as APOE2 and APOE3-Christchurch have also been reported to modulate lipid handling and AD vulnerability^16,17^. In addition, plasma ceramides decline with disease progression, whereas specific sphingomyelins and gangliosides show dynamic fluctuations potentially linked to synaptic remodeling^18,19^. Nevertheless, it remains unclear whether lipid dysregulation begins centrally and propagates peripherally, or whether peripheral lipid signals may themselves influence brain homeostasis.

This question is particularly compelling in autosomal dominant Alzheimer’s disease (ADAD), or Familial Alzheimer’s disease (FAD), where the Presenilin-1 (PSEN1)-E280A mutation provides a predictable model of disease evolution. Cross-sectional cohorts of this kind offer a unique opportunity to trace metabolic perturbations decades before clinical onset, establishing potential temporal patterns rather than correlation. In our Colombian kindred, both plasma p-tau217 and NfL discriminate PSEN1-E280A carriers from non-carriers in the early 20s, over two decades before symptom onset, while GFAP and YKL-40 show distinguishable patterns at different stages^1,2,20^. Yet, the contribution of lipid metabolism to this preclinical cascade remains poorly understood.

We hypothesize that lipidomic remodeling represents an early and dynamic marker of vulnerability in ADAD, reflecting the brain’s attempt to maintain structural and energetic balance under chronic metabolic stress. In this view, serum lipid profiles could serve as active indicators of brain homeostasis, enabling detection of preclinical alterations well before conventional protein biomarkers become abnormal.

To test this, we analyzed serum lipid profiles from 313 participants of the Colombian PSEN1-E280A cohort, double-blinded for genotype and stratified by age, sex, APOE isoforms, and cognitive status, using high-resolution mass spectrometry (HR-MS). Machine-learning models identified age-dependent lipid signatures discriminating carriers from non-carriers with high accuracy (AUC 8–93%). Latent Profile Analysis (LPA) further delineated five lipidomic endophenotypes, corresponding to progressive metabolic trajectories from childhood resilience to adult dysregulation. Integrating these peripheral findings with postmortem analyses of cingulate cortex and subventricular zone (SVZ) tissue, we uncovered coordinated changes in lipid enzymes and transporters, particularly sphingomyelinases, which suggest a bidirectional coupling between cerebral and systemic lipid homeostasis.

Altogether, this study addresses a critical gap in AD biology: the missing link between central lipid metabolism and its peripheral reflection during the earliest stages of disease. By reconstructing the developmental and metabolic trajectory of lipid dysregulation in FAD, we aim to establish lipidomic as a bridge between neurobiology and precision prevention, illuminating how peripheral lipid states encode cerebral vulnerability and how they may be harnessed to anticipate or modulate neurodegenerative risk.

## RESULTS

### Study design

We studied 313 participants from the Colombian PSEN1-E280A kindred with familial FAD (Supp. Table 1), including 175 mutation carriers (56%) and 138 non-carriers (44%), balanced by sex (55% female) (Fig 1b). Participants ranged from 6 to 72 years, they were predominantly children (6–12 years, 38%) and adolescents (13–19 years, 15%) (Fig. 1c). APOE genotyping showed predominance of ε3/ε3 (n = 204), followed by ε2/ε3 (n = 37) and ε3/ε4 (n = 55), with rare isoforms in the remaining participants. Considering all participants, among carriers, 22% exhibited cognitive impairment (mild cognitive impairment or dementia), while 90% were cognitively unimpaired and 2% reported subjective cognitive decline. Most individuals were free of metabolic (81%) and neuropsychiatric or neurological comorbidities (77%), and BMI (Body mass index) values were within the normal range (Supp. Table 1).

**Figure 1.**
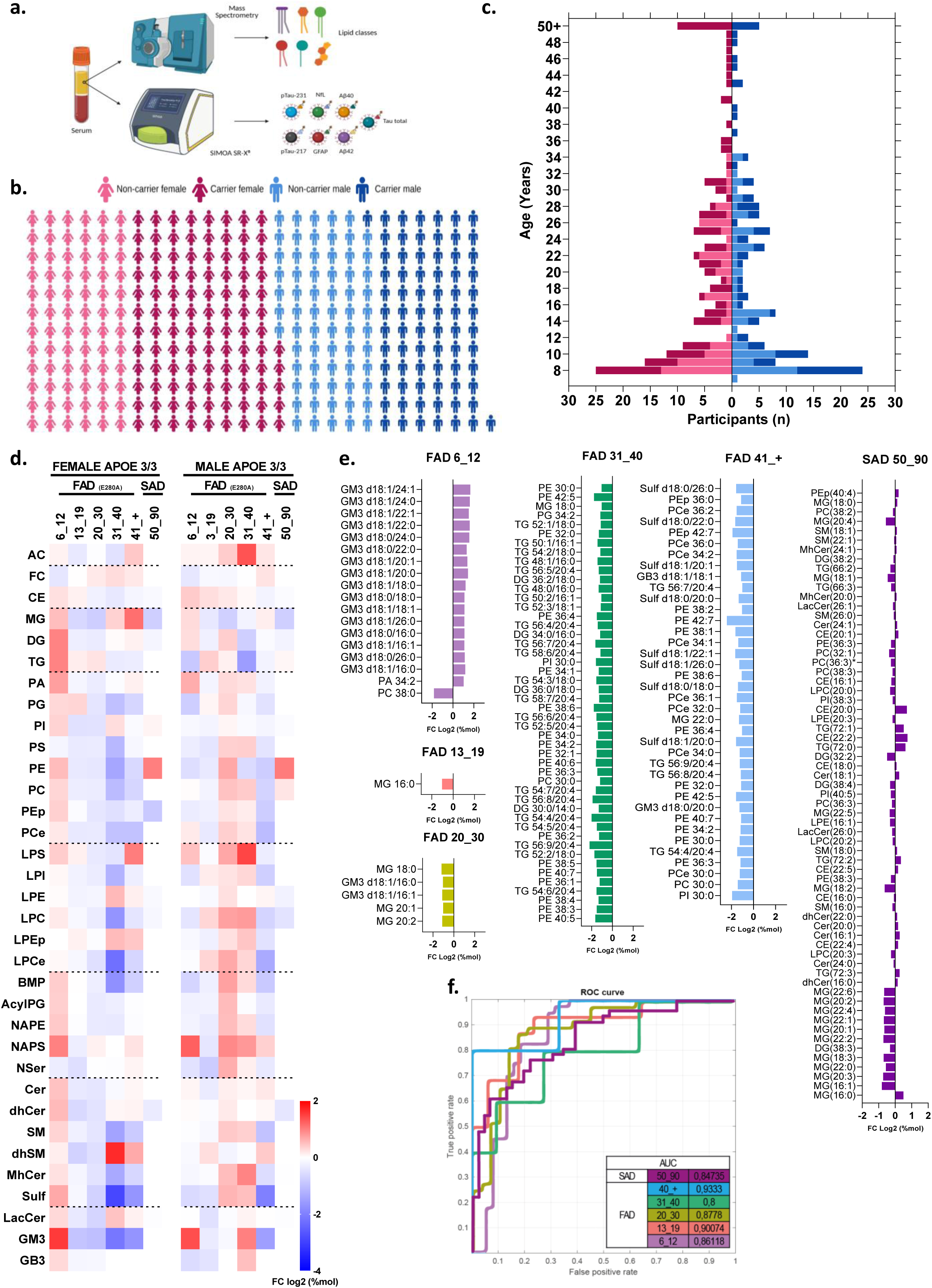
Age-dependent trajectories of the serum lipidome reveal early metabolic remodeling in PSEN1-E280A carriers (APOE 3/3 only). (a) Study design and serum sampling workflow for lipidomic (N=316) and SIMOA (N=173) protein analyses (created with BioRender.com). (b) Distribution of sex and PSEN1-E280A carrier status across the cohort. (c) Age distribution by sex (female, male) and PSEN1 status. (d) Heatmap showing log₂ fold-change (%mol) trajectories of lipid classes across age ranges (blue, negative; white, no change; red, positive), grouped by metabolic family: acyl-CoA derivatives (AC); sterols (free cholesterol [FC], cholesteryl esters [CE]); glycerides (mono-, di-, and triacylglycerols: MG, DG, TG); phosphoglycerides (PA, PG, PI, PS, PE, PC, PEp, PCE); lysophosphoglycerides (LPS, LPE, LPC, LPEp, LPCE); modified phosphoglycerides (BMP, acylPG, NAPE, NAPS, NSer); sphingolipids (Cer, dhCer, SM, dhSM, mhCer, Sulf); and glycosphingolipids (LacCer, GM3, GB3), in the SAD cohort, analyses were restricted to CE, MG, DG, TG, PI, PE, PC, LPE, LPC, Cer, dhCer, SM, mhCer, and LacCer species. (e) Age-dependent fold-change profiles of significantly altered species across childhood (6–12 y), adolescence (13–19 y), early adulthood (20–30 y), middle age (31-40 yo), FAD symptomatic stages (≥41 y), and SAD (50-90 y). Early metabolic divergence was marked by increases in GM3 and PA species during childhood, preceding broader declines in multiple lipid classes with age. (f) Receiver operating characteristic (ROC) curves for lipid-based classification models across FAD age groups and SAD (AUC = 0.8–0.93, accuracy ≥ 0.82).

### Lifespan trajectories of peripheral lipid remodeling in familial and sporadic Alzheimer’s disease

High-resolution lipidomic profiling quantified 593 lipid species across 34 classes to assess systemic alterations associated with the PSEN1-E280A mutation under an APOE ε3/ε3 background. Early lipid shifts were already evident during childhood (6–12 years), particularly in acylglycerols (MG, DG, TG), phospholipids, modified phospholipids, and glycosphingolipids such as GM3 and GB3 (Fig.1d). Sex-stratified analyses revealed distinct trajectories, with females showing transient early increase followed by decline in adulthood, and males displaying a more gradual elevations of neutral and sphingolipid species. But both sexes showed an inverse relationship between Free Cholesterol/Cholesteryl Ester (FC/CE); between SM (sphyngomyelin) and gangliosides GM3; and between GB3/dhSM (dihydro-SM).

Specifically, fold-change and significance analyses (%mol and p-values, Fig. 1e) identified GM3 as the most prominently exported lipid class into the bloodstream in children (6–12 years; however, an inflection occurs at 13-19 yo only showing a reduction in monoacylglycerol (MG) species, persisting into symptomatic stages (≥41 years). In addition, significant progressive reduction of another species as GM3 at 20 yo, TAG species and phospholipid (PC, PE) species, mainly at 31-40 and being more diverse including sulfatides and PL plasmalogens species in the older people. Also, a diverse profile in sporadic Alzheimer’s disease (SAD) was observed but exacerbating the reduction of different MG species.

Multivariate analyses by PCA (Principal component analysis) (Ext. Fig. 1a, b-b’) and PLS-DA (Ext. Fig. 1a, c-ć) captured global lipidome dynamics across age, revealing pronounced changes in specific lipid respect to abundance and discrimination respectively. In FAD, a notable shift in the FC/CE ratio was observed around 20 years of age, reflecting overall an inverse progressive abundance at the serum by PCA (Ext. Fig. 1a, b, b’); whereas GM3 species emerged as the most discriminant features by PLS-DA, showing a marked and progressive decline since 13 yo, alongside with a general inverse pattern of dhSM and NAPS species in all ages (Ext. Fig. 1a, c, c’). In SAD, although most MG decrease based on PCA, MG 18:0 is uniquely preserved and even increased compared to the others. Overall, among the lipid classes analyzed in SAD, MG species show the greatest variability, being predominantly the species rescued by PCA (Ext. Fig. 1a’, 1d–d’) and PLS-DA (Ext. Fig. 1a’, 1e–e’).

Overall, these results delineate a continuous trajectory of peripheral lipid remodeling, initiating childhood and dynamically progressing across the lifespan. SAD cases showed pronounced heterogeneity, yet both FAD and SAD cohorts ultimately converged on late-life lipidome deterioration. Critically, ROC analyses underscored the strong discriminative capacity of these lipidomic signatures across age groups, with AUC (Area under the curve) of 0.8–0.93, accuracy ≥0.82, and precision ≥0.80 (Fig. 1f), emphasizing their potential as early, robust bioindicator of metabolic divergence and neurodegenerative risk.

To visualize overall and temporal lipidome changes in PSEN1-E280A carriers versus non-carriers, including only ε3/ε3 isoform (Fig. 2a), or all isoforms (Ext. Fig. 2a); we generated a smoothed lineal regression using LOESS statistical approach, based on the average and standard deviation (SD) (Fig. 2b, Ext. Fig. 2a). Finding particularly relevant, an inflexion points of lipidome at 13-19 yo, which appear be inverted respect to 6-12 yo, with a more pronounced shift at the time in the carriers (Fig. 2c). Being the more impacting, the marked inversion of the ratios FC/CE, PE/TG, GM3/NAPS, dhSM/SM since 30 yo (Fig. 2 a). Lineal regression model depicted the inflexion point at 13-19 yo (Fig. 2b), highlighting the progressive increase of FC and decrease of CE, SM and MG in the PSEN1-E280A (APOE ε3/ε3) carriers (Fig. 2 b). With a sustained change of FC, CE and TAG by all isoforms (Ext. Fig. 2b). Interestingly, when we revised the levels of blood biomarkers we found a subtle trend of reduction of Aβ1-42, pTau231 and an increase of pTau217 mainly in females after 20 yo (Fig. 2 c-g), without notable changes in the other biomarkers and ratios in the carriers respect to non-carriers (Ext. Fig. 2c, g).

**Figure 2.**
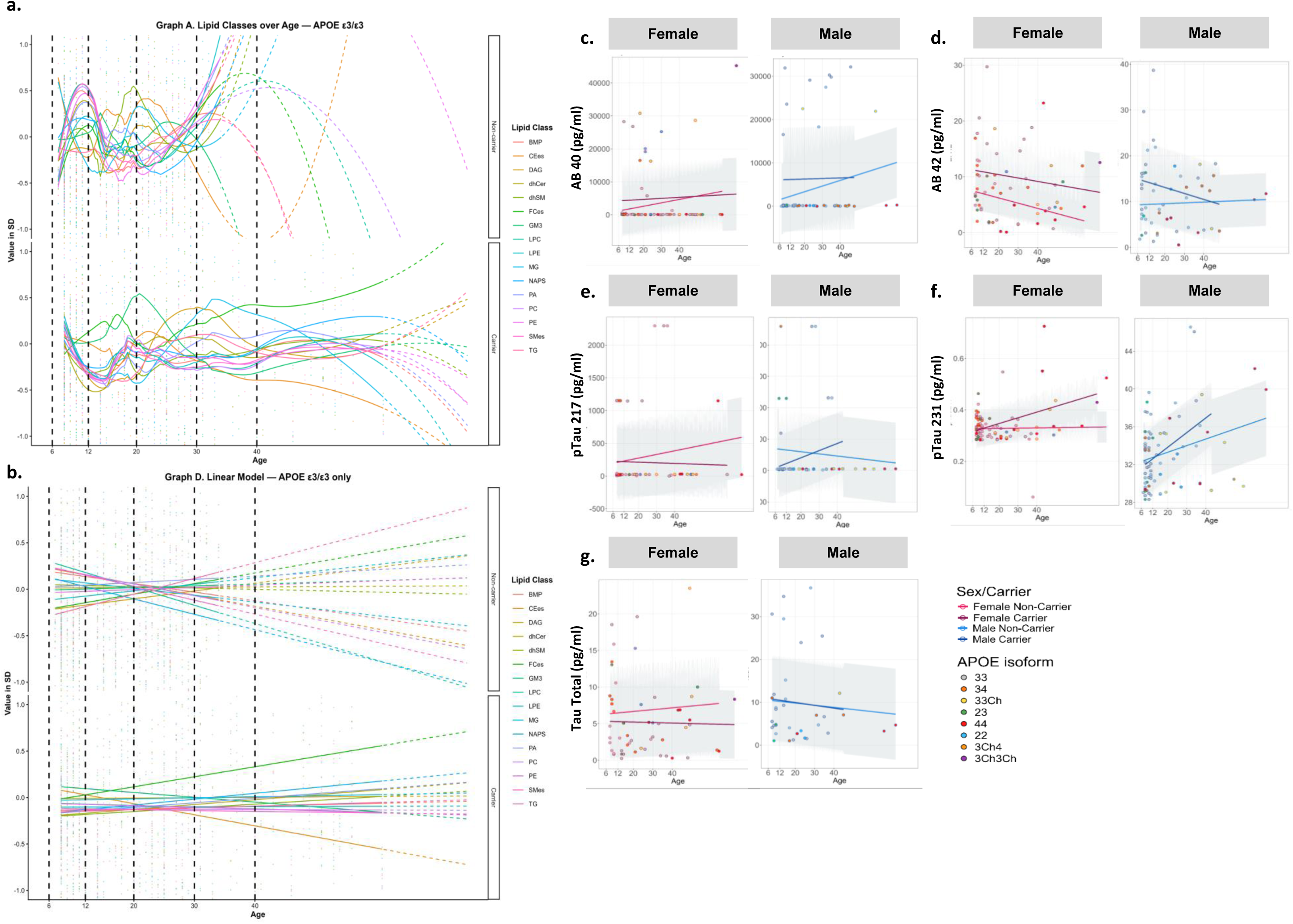
Age-dependent inflection points in lipidome remodeling and SIMOA biomarker trajectories. **(a)** Smoothed LOESS regression curves (mean ± SD) showing age-dependent trajectories of 16 representative lipid classes in PSEN1-E280A non-carriers with APOE3/3 (n= 207). **(b)** Corresponding linear-model fits for the same lipid classes (APOE3/3) (n=207). **(c–g)** SIMOA serum biomarker trajectories (pg/mL), displayed separately for females and males, and stratified by PSEN1-E280A carriers and non-carriers. including **(c)** Aβ40 (n=126), **(d)** Aβ42 (n=116), **(e)** pTau231 (n=126), **(f)** pTau217 (n=173), and **(g)** total tau (n=93). All SIMOA biomarkers (Aβ40, Aβ42, NfL, GFAP, pTau231, pTau217, and total tau) were analyzed across age using all APOE isoforms represented in the lipidomics subset.

### APOE isoform and sex-dependent modulation of peripheral lipid trajectories in PSEN1-E280A carriers

We next investigated whether coinheritance of APOE variants modifies the lipidomic trajectory of PSEN1-E280A carriers. While the cohort was predominantly homozygous for the wild-type allele (ε3/ε3; 61.3% of females, 70% of males), with rare occurrences of ε2/ε2 and ε4/ε4 genotypes (<3%; Ext. Data Fig. 3a, b), stratification by isoform revealed striking sex-specific metabolic nuances. Heatmap analysis demonstrated that lipid perturbations were markedly more pronounced and structured in females than in males (Ext. Data Fig. 3c). In young females (6–12 years), the presence of the ε2 allele (ε2/ε3) appeared to mitigate the early metabolic surge observed in ε3/ε3 carriers, dampening the elevation of acylglycerols (MG, TG) and phospholipids (PE). This relative lipid stability in ε2/ε3 females persisted through adolescence (13–19 years), contrasting with the increased concentration of these lipids seen in ε3/ε3 and ε3/ε4 genotypes.

**Figure 3.**
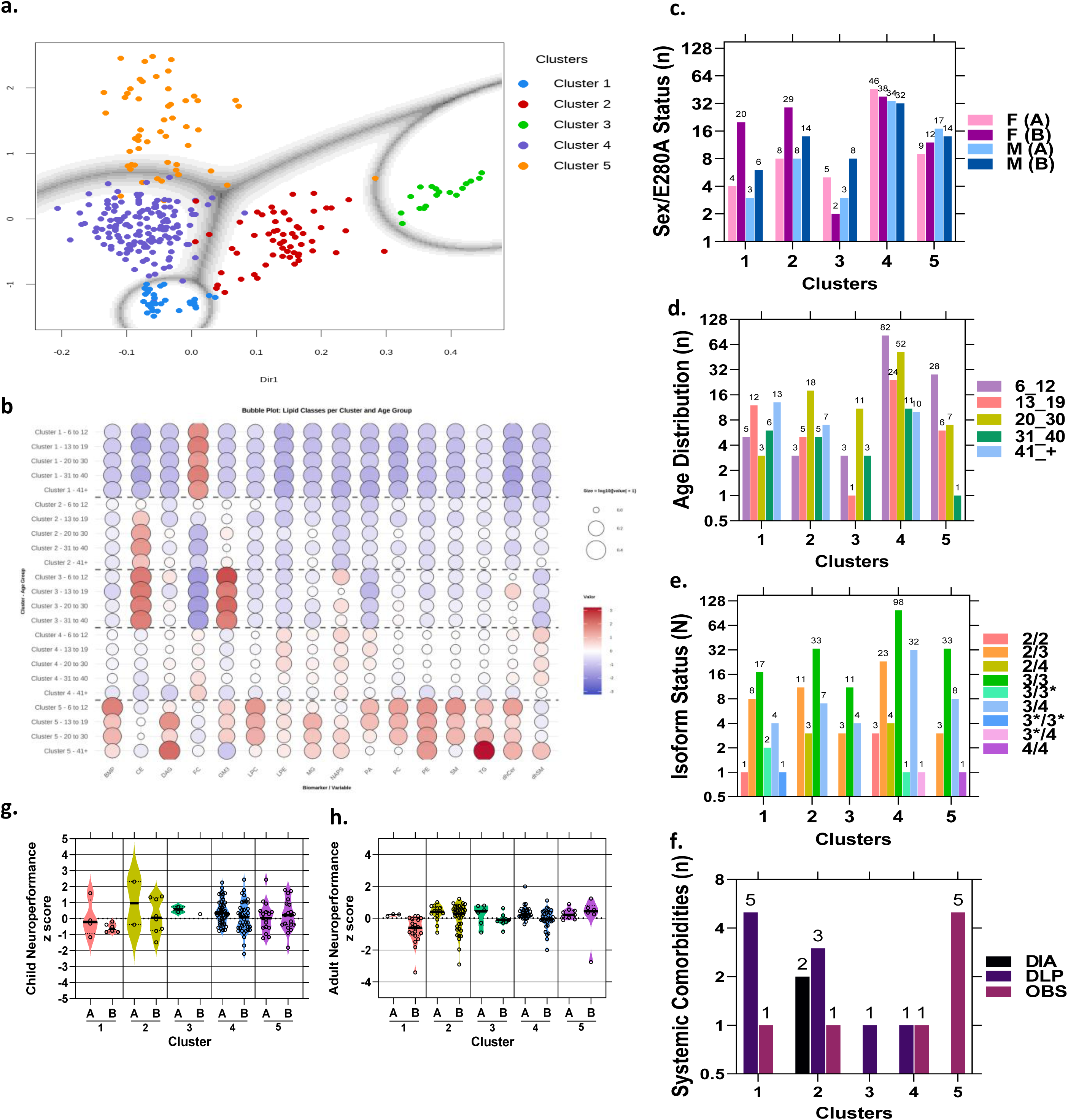
Latent lipidomic organization defines developmental and clinical endophenotypes aligned with lifespan inflection points. **(a)** Latent Profile Analysis (LPA) identifying five lipidomic clusters derived from 16 representative lipid classes. **(b)** Bubble plot illustrating inter-cluster similarity based on lipid class correlation. **(c)** Distribution of PSEN1-E280A non-carriers (A) and carriers (B), stratified by sex, across the five lipid-defined clusters. **(d)** Age distribution of participants within each lipidomic cluster. **(e)** APOE isoform distribution across clusters. APOE Christchurch variants are depicted with * along the allele **(f)** Distribution of metabolic comorbidities across lipid-defined clusters, including diabetes (DIA), dyslipidemia (DLP), and obesity (OBS). **(g-h)** Neuropsychological performance per cluster. **(g)** Children and adolescents (6–16 years) were evaluated using WISC-IV composite indices, while **(h)** adults (≥18 years) were assessed using the MMSE, CERAD’s Word List Recall, and the Semantic Verbal Fluency tests. All cognitive measures were normalized to z-scores and integrated into a global neuroperformance index to enable direct comparisons across age groups, with panels A and B representing non-carriers and E280A carriers, respectively.

Conversely, in adulthood (>30 years), the ε4 allele in PSEN1-E280A carriers was associated with a distinct risk signature, characterized by elevated MG and dhSM). Furthermore, older female carriers homozygous for APOE Christchurch variant (ε3*/ε3*) displayed specific increases in FC and MG (Ext. Fig 3c). In contrast to the dynamic isoform-dependent patterns observed in females, lipid trajectories in males lacked clear stratification by APOE genotype, suggesting a sex-dimorphic vulnerability to lipid dyshomeostasis in the preclinical phase of AD (Ext. Fig 3c).

### Latent profile analysis of lipid classes discriminates endophenotypes associated to clinical and neuropsychological characteristics

Latent Profile Analysis (LPA) based in lipidome of 16 lipid classes blinded for clinical stratification, identified five distinct endophenotypic clusters with common lipid profile (Fig. 3a–b). Cluster 1 showed elevated free cholesterol (FC, 1.69-fold), Cluster 2 increased cholesteryl esters (CE, 1.39), Cluster 3 combined high CE (2.0) and GM3 (2.3) with reduced FC (–1.75), Cluster 4 was metabolically neutral, and Cluster 5 exhibited broad enrichment in phospho– and sphingolipids—including PE (1.75), SM (1.86), BMP (2.05), LPC (1.99), GM3 (0.93), and PC (1.06)—representing the most metabolically divergent profile (Fig. 3b).

Demographic and genetic patterns (Fig. 3c–e) clusterized 1–2 enriched in PSEN1-E280A carriers (n = 69), predominantly female (n = 49), and included older adults (≥41 yo), Cluster 3, largely composed by male young adults (20-30 yo), Clusters 4–5 comprised younger participants (6–30 yo), mainly childrens (6-12 yo). APOE genotype ε3ε3, ε2ε3, ε3ε4 alleles were distributed in all clusters; ε2ε2 and 3Ch homo or 3 heterozygous in cluster 1, while ε4 homozygous and Christchurch heterozygous variant were present in Clusters 4-5 (Fig. 3e).

Also, lipidomic profiles tracked clinical and cognitive outcomes (Fig. 3f–h, Ext. Fig 4)). Metabolic comorbidities showed distinct distributions: obesity was more frequent in Cluster 5, whereas dyslipidemia and diabetes predominated in older adult clusters (1–2). In the global cognitive analysis, Cluster 1 showed the lowest performance for children and adolescents (Fig. 3g) and adults (Fig. 3h). When stratifying by carrier status, PSEN1-E280A carriers performed worse than non-carriers across all clusters, except in Cluster 5. Overall, the distribution of scores across individual neuropsychological tests were consistent with that pattern (Ext. Figure 4). Interestingly, Cluster 1 presented the highest levels of NFL, GFAP, and pTau217, and lowest for Aβ1-42 and pTau231, in the cohort of cases that were evaluated by SIMOA (Single molecule array) biofluid markers (Supp. Table 2; Supp. Fig. 2).

**Figure 4.**
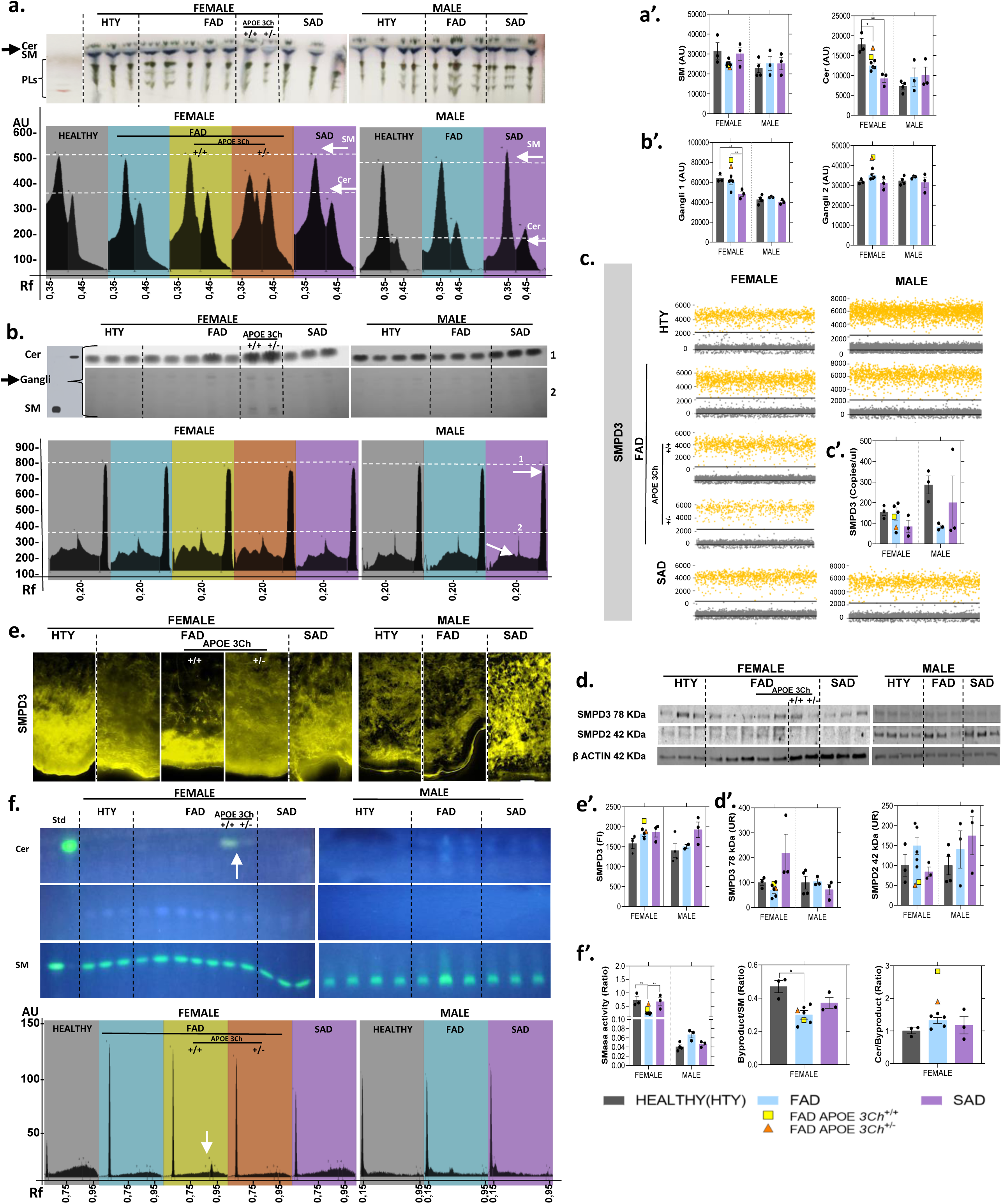
Sphingolipid metabolism and enzymatic regulation in postmortem cingulate cortex. **(a-b)** Representative thin-layer chromatography (TLC) plate and scan showing the separation of major sphingolipid classes, including sphingomyelin (SM), ceramide (Cer) **(a)**, and ganglioside-enriched fractions **(b)**. **(a’–b′)** Corresponding quantifications from TLC. **(c)** Representative scatter plots of individual dPCR reactions acquired using the ThermoFisher Q Studio dPCR platform, showing detection of SMPD3 mRNA in post-mortem cingulate tissue from women (left panel) and men (right panel). Each yellow dot corresponds to a positive droplet, defined by fluorescence above the automated threshold, whereas grey dots represent negative droplets. Individuals are grouped as healthy controls (HTY), carriers of FAD–PSEN1 E280A, carriers of FAD–PSEN1 E280A with the protective APOE3Ch+/+ variant, simple heterozygotes, and cases of sporadic Alzheimer’s disease (SAD). **(c′)** Bar plot showing the absolute quantification of SMPD3 (copies/µL) across diagnostic groups. Quantification of *smpd3* expression by digital dPCR. **(d–d′)** Quantification of SMPD2 and SMPD3 protein levels by western blot. **(e–e′)** Representative immunofluorescence (IF) images showing the cortical distribution of SMPD3 and corresponding fluorescence quantifications. **(f)** Representative enzymatic activity assay of sphingomyelinase (SMase) a scan **(f’)** and corresponding quantifications. Data are expressed as mean ± SEM; significance: *p < 0.05, **p < 0.01, ***p < 0.001 (N=23). For women, groups include healthy controls (HTY, n=), FAD–PSEN1 E280A carriers (n=5), carriers with the protective APOE3Ch+/+ variant (n=1), APOE3Ch+/– heterozygous carriers (n=1), and sporadic AD cases (SAD, n=3). For men, groups include HTY (n = 4), FAD–PSEN1 E280A (n=3), and SAD (n=3). Parametric comparisons were performed using one-way ANOVA, whereas non-parametric data were analyzed using the Kruskal–Wallis test.”

### Lipid metabolic changes in postmortem cingulate cortex from Alzheimer’s disease brains

Guided by systemic lipid alterations in familial and sporadic AD, we assessed key lipid enzymes and transporters in the cingulate cortex and contiguous areas as corpus callosum, and SVZ through several analitical techniques (Ext. Fig. 5a). Protein quantification revealed selective, sex and disease condition-dependent remodeling. Female SAD showed marked upregulation of ACSL4, PLTP, and Erlin2, while FAD femaleswere influenced by APOE3 Christchurch dosage increasing SREBP2 and MFSD2A. Gene Ontology analysis supported enrichment of SREBP signaling, cholesterol and lipid biosynthesis, and lipid transport pathways (FDR < 0.005), linking systemic lipid signatures to central biochemical remodeling and implying downstream alterations in cortical lipid composition and organization (Ext. Data Fig. 5 a–c).

**Figure 5.**
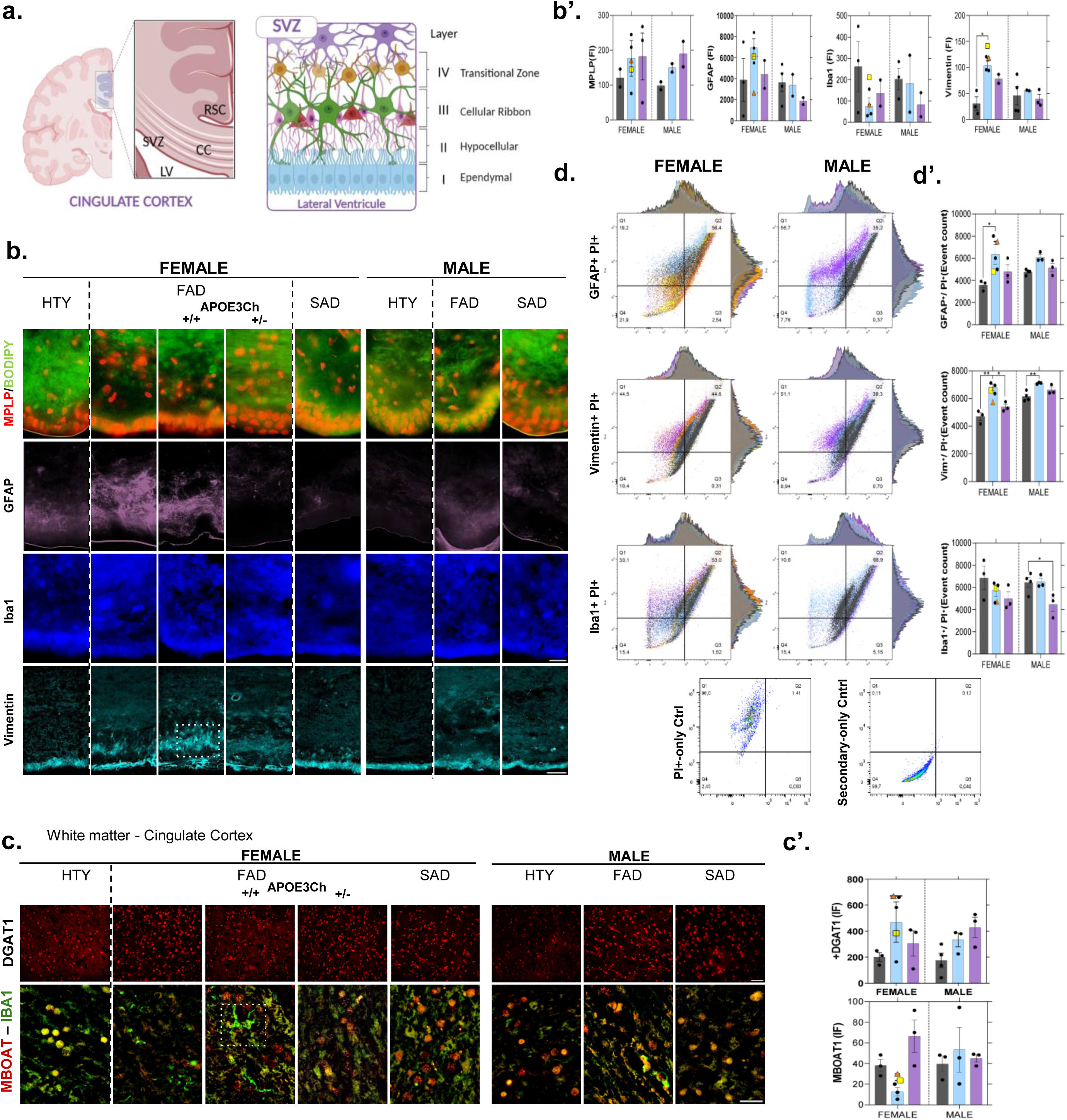
Glial remodeling in postmortem cingulate cortex and adjacent regions. **(a)** Representative schematic of the cingulate cortex, corpus callosum, and subventricular zone (SVZ), showing their anatomical location (left) and laminar organization (right). Layers are defined as: I, ependymal; II, hypocellular; III, cellular ribbon; IV, transition zone. **(b-b’)** Representative immunofluorescence (IF) images showing MPLP (red), BODIPY (green), GFAP (magenta), IBA1 (blue), and Vimentin (cyan) labeling across groups and genotypes, used for glial characterization of the SVZ, together with the corresponding fluorescence quantifications. MPLP, BODIPY, GFAP, and IBA1 panels were acquired at 60X magnification (scale bar=20 µm). Vimentin and DGAT1 at 20X magnification (scale bar=50 µm). **(c–c′)** IF staining of DGAT1 (red), MBOAT1 (red), and IBA1 (green) in the cingulate cortex and corpus callosum, illustrating molecular and cellular alterations across conditions. **(d–d′)** Flow cytometry analysis and corresponding quantifications of GFAP⁺, Vimentin⁺, and IBA1⁺ populations. Data are expressed as mean ± SEM; significance: *p < 0.05, **p < 0.01, ***p < 0.001 (N=23). For women, groups include healthy controls (HTY, n= 3), FAD–PSEN1 E280A carriers (n=5), carriers with the protective FAD APOE3Ch+/+ variant (n=1), FAD APOE3Ch+/– heterozygous carriers (n=1), and sporadic AD cases (SAD, n=2-3). For men, groups include HTY (n = 2-4), FAD–PSEN1 E280A (n=2-3), and SAD (n=2-3). Parametric comparisons were performed using one-way ANOVA, whereas non-parametric data were analyzed using the Kruskal–Wallis test.”

In this same region, we profiled neutral and polar lipid classes, including phospholipids and sphingolipids (Ext. Data Figs. 6, 7). Neutral lipids showed subtle but organized reorganization: FC and CE were largely unchanged, though APOE3Ch (APOE Christchurch variant) carriers exhibited a consistent downward trend in the accumulation of these lipids measured by TLC assay (Ext. Data Figs. 6a, b). DGAT2 gene had a reduced expression in FAD and SAD males, and Triglycerides displayed an increase in FAD females and SAD males, supported by DGAT1 protein levels, without changes by IF increase in DAGT1 and ABCA1 (Ext. Data Fig. 6 b, b’-6 e-é). Interestingly, DGAT1 IF in FAD females independent of APOE isoform revealed a loss of cell stratification in SVZ (Ext. Data Fig. 6e).

**Figure 6.**
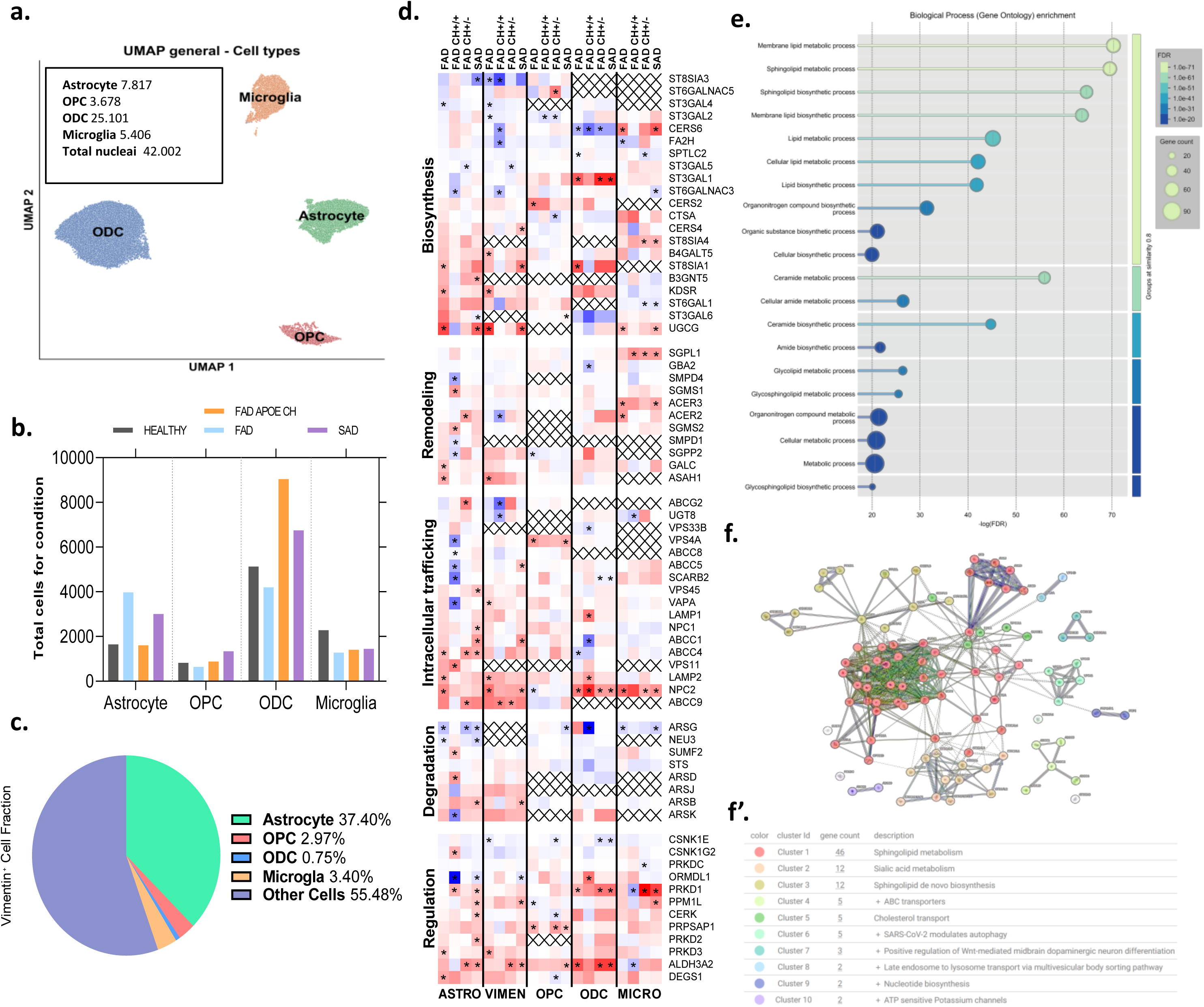
Single-nucleus transcriptomic profiling of sphingolipid metabolism in glial populations. **(a)** UMAP visualization of 31,919 nuclei from control, FAD, APOECh, and SAD samples, grouped into four major clusters: oligodendrocytes (ODC), oligodendrocyte precursor cells (OPC), astrocytes (Astro), and microglia (Micro). (b) Distribution of glial cell types according to Alzheimer’s disease diagnosis. **(c)** Distribution of Vimentin expression across glial populations. **(d)** Heat map fold change analysis of differentially expressed genes from single-cell data (blue, negative; white, no change; red, positive); statistical significance indicated by asterisks (*p < 0.05). **(e)** Gene Ontology (GO) enrichment analysis showing biological processes associated with the differentially expressed genes. **(f)** Clustering of genes based on co-expression patterns. **(f′)** Functional annotation of gene clusters (N = 12). In this analysis, samples were not stratified by sex and included healthy controls (HTY, n = 3), FAD–PSEN1 E280A carriers (n = 3), APOE3Ch+/+ carriers (n = 1), APOE3Ch+/– carriers (n = 2), and sporadic AD cases (SAD, n = 3).

In the case of phospholipids, as PS, PE, and PC, were depleted in FAD and SAD females, with a more significance in SAD ones (Ext. Data Figs. 7a, b, b**’**). p-cPLA2 showed an increased in FAD and SAD, but LPCAT1 and MBOAT1 enzymes did not change; however, the ε3Ch/ε3Ch woman case, presented a reduction of p-cPLA2 and increase at the acyltranferases (Ext. Data Fig. 7c, c**’**). Particularly, LPCAT1 IF presented a differential pattern of layering at the layer I and II of the SVZ in all FAD females (Ext. Data Fig. 7d, d**’**) and MBOAT1^+^ cells had more dispersion in SAD conditions. All these proteins confirm a crucial role in phospholipid remodeling by gene ontology (Ext. Data Fig. 7e).

Respect to sphingolipids, SM and ceramide classes were reduced in both female FAD and SAD cortices relative to controls, especially Cer in SAD (Fig. 4a, a’); while APOE3Ch carriers maintained relatively higher Cer levels. Ganglioside-enriched fractions revealed polarity-dependent shifts, with the less polar fraction (Gangli 1) and more polar fraction (Gangli 2), were upregulated in APOE3Ch carriers (Fig. 4b, b’). At the molecular level, dPCR detected no overall SMPD expression changes (Fig. 4c); whereas protein analysis revealed an elevation of SMPD3 and a reduction of SMPD2 in FAD APOE3Chs (Fig. 4d –d’). SMPD3 IF confirmed an enhanced SMPD3 signal in FAD APOE3Ch with an increased distribution in layer II of the SVZ (Fig. 4e). Complementarily, in situ sphingomyelinase (SMase) activity was elevated in FAD APOE3Chs, aligning with preserved Cer pools and enhanced turnover (Fig. 4f–f’, arrow).

### Glial characterization and transcriptional expression of sphingolipid pathway genes in postmortem Alzheimer’s disease brains

To determine whether lipid remodeling could be associated with structural and transcriptional glial alterations. First, we analyze and focus on main changes observed in white matter (WM) and SVZ of postmortem brains from familial FAD, SAD, compared with healthy controls (HTY). Immunofluorescence revealed an increased redistribution of glial cells, mainly Oligodendrocytes (MPLP+), Astrocytes (Vimentin+ and GFAP+) at layers I and II of the SVZ in FAD, especially in females and a particular reduction of microglia (Iba1) in SAD (Fig. 5a, b, b’). Also, we found more DGAT1 positive cells in white matter of female FAD (Fig. 5c, c’). Changes on cell morphology of Vimentin+ and Iba1+ cells were observed under APOE3Ch homozygous with a more expanded phenotype for astrocytes and microglia, respectively (Fig. 5b). Flow cytometry confirmed an increase in GFAP⁺ and Vimentin⁺ populations, predominantly in FAD, accompanied by a reduction in Iba1⁺ cells in SAD (Fig. 5 d, d’). Importantly, cell-cycle analysis by propidium iodide (PI) staining revealed increased cell-cycle activity across all phases (G1, S, and G2/M), with the most prominent shifts occurring in Vimentin⁺ populations, particularly in females with FAD (Supp. Fig. 3).

To understand the meaning of sphingolipids as main actors in brain and peripheral lipidome, and glial cells rescue as the main morphology changing in FAD and APOE3Ch+/+; we use Single-nucleus RNA sequencing delineated by cell-type-specific expression of genes in the prefrontal cortex from the postmortem GNÁs bank available in (GEO accessions: GSE222494, GSE222495, GSE206744). We found a marked differentiated oligodendrocytes (ODCs), followed by astrocytes, microglia, and oligodendrocyte precursor cells (OPCs) (Fig. 6a). Being more abundant ODC in all AD conditions, mainly in FAD-APOE3Ch and astrocytes in FAD cases (Fig. 6b). Because, Vimentin had an interesting morphological shift by previous IF in glial ribbon at the SVZ (Fig. 5b); vimentin+ cell expression was revised in all glial population, and an increase in astrocytes was found in all samples, providing a possible link between vimentin + cells and astrocyte morphology in the IF analysis (Fig. 6c, 5b).

Gene-level analyses categorized by sphingolipid-related genes according to biosynthesis, remodeling, intracellular trafficking, degradation, and regulation showed a notable down regulation of the genes in the APOE3Ch condition, mainly in vimentin+ cells and ODC and downregulated immune response and degradation in microglia (Fig 6d). Supported by Gene Ontology (GO) enrichment (Fig. 6e) and clustering analyses (Fig. 6f-f’) which highlighted the predominant cellular functions associated to sphingolipid metabolism, ceramide and sialic acid metabolism.

## Discussion

Our study identifies, for the first time, a childhood serum lipid signature in PSEN1-E280A mutation carriers that undergoes inversion during adolescence. This trajectory suggests an early failure in the clearance of proinflammatory lipids, which intensifies in adulthood alongside rising p-tau217, disrupted ganglioside metabolism, and increased vulnerability in females and APOE4 carriers. APOE3^Christchurch carriers, by contrast, displayed distinct lipid salvage profiles, highlighting genotype-specific resilience. The early lipid imbalance, characterized by reduced CE influx and local lipid synthesis, appears to activate ceramide salvage in glial populations, suggesting a compensatory adaptation. At the same time, reduced oligodendrocyte pruning and downregulation of lipid-related genes in microglia (e.g., *prkd1, ppml1, aldh3a2, ugt8*) reflect impaired immune engagement. These metabolic changes are different in sporadic AD (SAD), where alterations instead align with systemic stress and proinflammation.

Although AD spans both genetic and sporadic forms, our findings suggest that lipid metabolism may represent a shared, upstream vulnerability. In PSEN1-E280A carriers, distinct lipidomic alterations, particularly involving cholesterol, sphingolipids, and glycolipids, emerge as early as 6–12 years of age. Elevated GM3, decreased FC, and increased CE in preadolescence mark a genotype-driven reprogramming of systemic lipid homeostasis. These findings indicate an early breakdown in the sphingoglycolipid–cholesterol axis that underpins neuronal membrane integrity and synaptic architecture. Given the CNS enrichment of complex gangliosides, the elevated serum GM3 levels likely reflect impaired neural retention or enhanced efflux, consistent with early lipid trafficking defects^21–23^.

Experimental models underscore the role of ganglioside metabolism in early neurodevelopmental processes. Genetic disruption of enzymes such as *St3gal5 or B4galnt1* impairs myelin formation^24^, while abnormal myelination precedes amyloid or tau pathology and is linked to microglial activation in the 3x-Tg-AD model^25^. These structural changes are mechanistically coupled to APP processing^26^. Positioning ganglioside metabolism as a central vulnerability pathway bridging neurodevelopment and neurodegeneration.

In non-carriers, childhood GM3 decline and transient pTau217 increases likely reflect biological processes of synaptic pruning and lipid turnover during brain maturation^27^. By contrast, PSEN1-E280A carriers exhibit persistent GM3 elevation, a sustained imbalance in the CE:FC ratio, and reduction of MG 16:0 beyond adolescence, indicating early divergence from healthy lipid developmental trajectories and preceding conventional biomarker conversion^28^.

This metabolic shift deepens in adolescence and early adulthood, marked by reductions in CE:FC, GM3, and MG species, inversion of dhSM/SM, and sustained N-acylphosphatidylserine (NAPS) elevation. NAPS are implicated in membrane repair and neuroprotection (Guan et al., 2007; L. Wood, 2015)^29^, and their rise suggests compensatory membrane remodeling. These changes are more prominent in females, aligning with sex-linked differences in lipid homeostasis and myelination^30,31^.

By age 31–40, lipid dysregulation becomes more pronounced, with declines in phosphatidylethanolamines (PE), phosphatidylglycerols (PG), gangliosides, DAG, and TAG. Meanwhile, dhSM, NAPS, and N-acylserines (NSer) are enriched, consistent with stress-adaptive remodeling under inflammatory load. This is accompanied by reduced Aβ1-42 and increased pTau217 (Fig. 2), suggesting a transition from developmental plasticity to a metabolically constrained state, coinciding with early PET-detectable amyloid/tau accumulation^32–34^.

Unlike protein biomarkers, which reflect threshold-crossing events, lipidomics provides a continuous and mechanistically informative readout of presymptomatic processes. Their temporal structure, emergence in childhood, and alignment between brain and periphery point to lipids as early biomarkersof metabolic fragility, supporting their utility for preventive therapeutics.

An unsupervised Latent profile analysis (LPA) further revealed that systemic lipid organization reflects developmental, genetic, and cognitive phenotypes. Cluster 4, the most abundant, captured children and adolescents with gradual FC and dhSM release. Cluster 5, less extensive, showed broad lipid export, particularly DAG, TAG, MG, and PE. This pattern may represent a transient adaptive remodeling, possibly reminiscent of cognitively resilient or presymptomatic ADAD individuals^35,36^.

Clusters 1 and 2, dominated by older individuals, showed elevated FC or CE and diminished lipid clearance. Cluster 3, comprising mostly young adults, revealed intermediate susceptibility, defined by CE and GM3 release. Cognitive performance declined progressively from Cluster 5 to Cluster 1, with Cluster 1, composed largely of female carriers with dementia, showing the lowest Neuropsychological scores. Interestingly, early lipid profiles also aligned with clinical comorbidities, including dyslipidemia and diabetes (Clusters 1–2), and obesity (Cluster 5), consistent with earlier findings of “compensatory” metabolic phenotypes in preclinical AD^37,38^. APOE4 carriers presented elevated FC irrespective of age or sex but did not cluster uniquely, except for a 6–12-year-old APOE4/4 PSEN1-E280A boy placed in Cluster 5.

The presence of the APOE3Christchurch (APOE3Ch) allele, linked to delayed onset, was observed in both metabolically altered and resilient clusters. These results suggest that APOE3Ch may buffer, but not eliminate, lipid-mediated vulnerability, in line with its reported context-dependency^16^.

Collectively, this latent lipidomic architecture supports a model in which systemic metabolism during development encodes vulnerability gradients for future neurodegeneration. Beyond serving as biomarkers, these profiles map inflection points where metabolic flexibility wanes and pathogenic susceptibility increase.

We next investigated whether these peripheral lipid trajectories mirrored central remodeling. Postmortem brain samples revealed sex– and genotype-dependent expression of lipid regulators. SAD females, and to a lesser extent FAD female, showed increased ACSL4, PLTP, and ERLIN2, along with TAG accumulation and proinflammatory p-cPLA2 signaling, suggestive of ferroptosis-associated lipotoxicity^39,40^. In contrast, APOE3^Ch homozygotes exhibited upregulation of SREBP2, MFSD2A, LPCAT1, and MBOAT1, consistent with a compensatory response to diminished CE, FC, and structural phospholipids in the brain parenchyma^41,42^.

Sphingolipids (SL) were globally reduced in both FAD and SAD cortices yet relatively preserved in APOE3^Ch carriers. Enhanced sphingomyelinase (SMase) activity in these cases supports a functional salvage pathway^43,44^. In contrast, classic FAD cases showed accumulation of intermediate lipid species like Cer-1P, consistent with elevated cellular stress.

Glial profiling further revealed increased DGAT1, MPLP+, and GFAP+/Vimentin+ cells in female FAD brains. SVZ-specific Vimentin+ increases in APOE3^Ch carriers aligned with transcriptomic evidence of microglial lipid metabolism downregulation (*prkd1, aldh3a2, ugt8*). suggest an immune evasion mechanism that might also impact lipid metabolism, as has been shown that microglia can impact the accumulation of lipid species within the brain^45^. Nonetheless, deeper mechanism studies are warranted.

Together, these results outline a multiscale model in which peripheral lipid remodeling reflects, and possibly anticipates, central cellular vulnerability. Lipid dysregulation emerges not merely as a downstream consequence of proteinopathy, but as a co-initiating or amplifying process involving glial, metabolic, and neurodevelopmental pathways.

Sex and genotype stratification of lipid profiles offers a refined lens on disease risk, particularly during developmental stages when metabolic plasticity is highest. These insights support the use of lipidomics to enhance predictive models and identify modifiable targets.

The identification of the subventricular zone (SVZ) as a site of structural-metabolic coupling further highlights its potential as an intervention point. Modulation of acyltransferases, lipid transporters, or glial support pathways may offer therapeutic avenues to preserve lipid homeostasis and delay onset.

In conclusion, this work supports a conceptual shift in Alzheimer’s research, from a late-stage proteinopathy to an early, lipid-centered framework. Lipids function not only as biomarkers but also as modulators of neurodegenerative risk. Integrating lipidomic, transcriptomic, and imaging data across the lifespan will be essential to determine whether rebalancing brain–periphery lipid coupling can alter Alzheimer’s disease trajectories in both familial and sporadic forms.

## Data availability

Mass spectrometry–based lipidomics data have been deposited in the Nbiol-GNA-UdeA/E280A-Serum-Lipidomics-Repository-GNA-Cohort: E280A Serum partner repository (dataset identifier: [![DOI](https://zenodo.org/records/17693328)] (https://doi.org/10.5281/zenodo.17693327). This repository contains serum lipidomic profiles from 316 individuals of the Colombian PSEN1-E280A kindred, curated by the Grupo de Neurociencias de Antioquia (GNA), spanning ages 6–≥41 years and including APOE genotypes. All processed data supporting the findings of this study are available in the Supplementary Information and Source Data files. All reagents used in this study are commercially available, and catalog numbers are listed in the Reagents Table (Supp. Table 4 and 5). Source data are provided with this paper.

## Supporting information

Supplementary comprehensive reagents

## Acknowledgements

This work was supported by Minister of Science and Technology Colombia (Minciencias-ICETEX, code #82336 to GPCG), National Institute on Aging and CODI-University of Antioquia (to GPCG), the U.S. National Institutes of Health (NIA, R21AG079574 to GPCG; R01-AG056387-01 to EA-G; R01NS117538 to EA-S), the Spanish Ministry of Science, Innovation and Universities (PID2021-126818NB-I00 to EA-G). Columbia University by Metabolon SAD data set. We thank Renu Nandakumar for assistance with the lipidomics analysis.

## Author Contributions Statement

Conceived the project: GPCG and EA-G. Designed experiments: GPCG, IDSC, DCAA, JDAL and EA-G. Generated experimental data: IDS-C, LALT, NDGG, SCAC, JPBC. Collected patient samples: EH, JPBC, AV, DFAN, FL. Collected/analyzed lipidomics data: GPCG, IDSC, NGG, MHR, LMTCh, GJF, EO, YQ, ES, DCAA, JAAL and EA-G. Wrote the manuscript: GPC-G, IDSC, EA-G. Critically edited the manuscript: all authors. Approved final version of the manuscript: all authors.

## Figure legends

**Extended Data Fig. 1.**
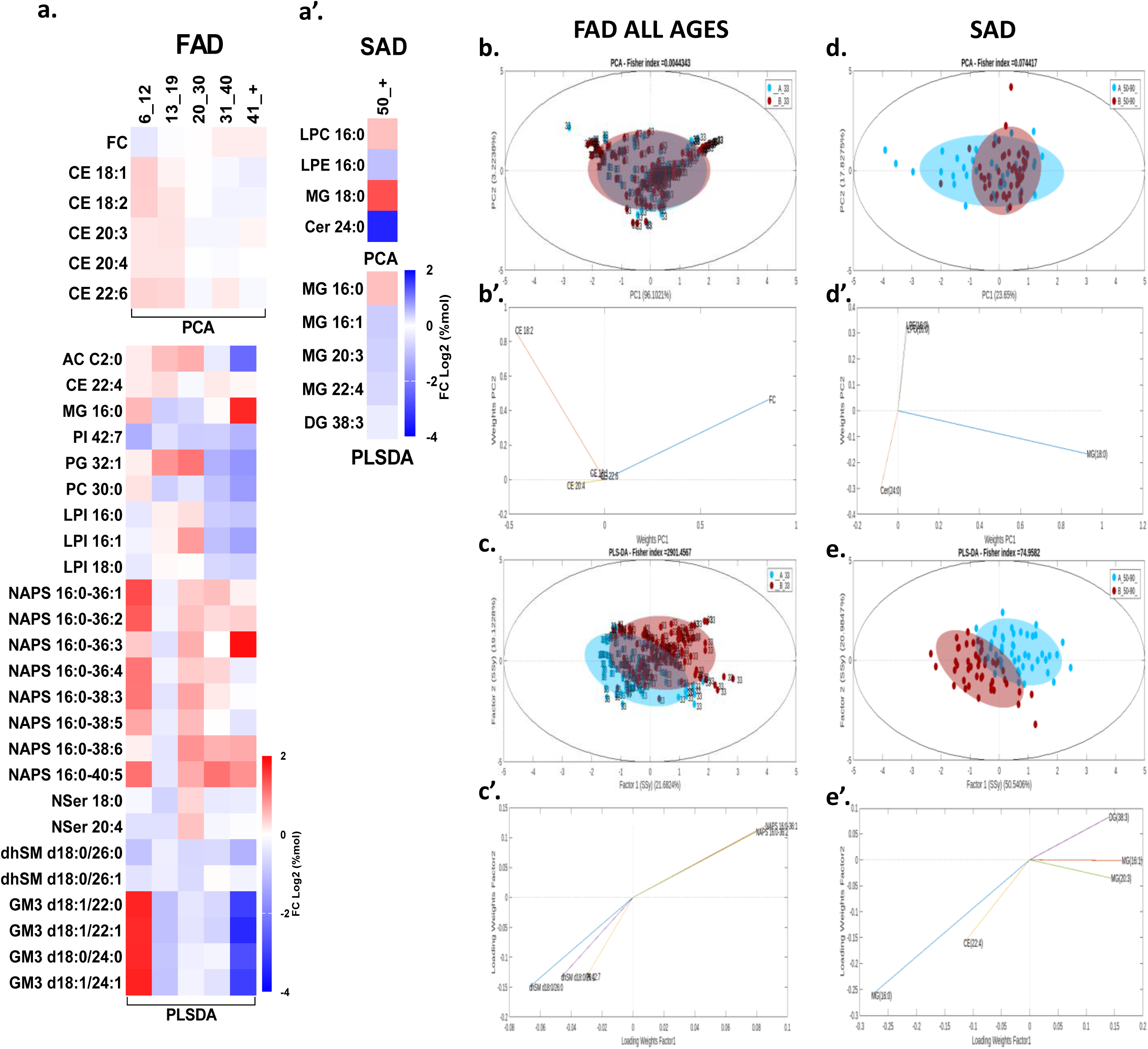
Multivariate structure and discriminant lipid species underlying PSEN1-E280A age trajectories and SAD group differences. **(a)** Heatmap of discriminant lipid species derived from PCA and PLS-DA models for FAD, built using lipid species that significantly discriminated groups at each age point, and grouped to visualize their temporal trajectories across childhood, adolescence, and adulthood (blue, negative; white, no change; red, positive). **(a’)** Heatmap of discriminant lipid species derived from PCA and PLS-DA models for SAD (blue, negative; white, no change; red, positive) **(b,b′; d,d′)** Graphical PCA of the full dataset (n = 313), integrating all ages and both sexes, to capture the global lipidome variance structure by age and carrier status in FAD **(b,b′)** and SAD **(d,d′). (c,c′; e,e′)** Graphical PLS-DA including all ages and both sexes, illustrating separation between PSEN1-E280A mutation carriers and non-carriers in FAD **(c,c′)** and between patients and controls in SAD **(e,e′).**

**Extended Data Fig. 2.**
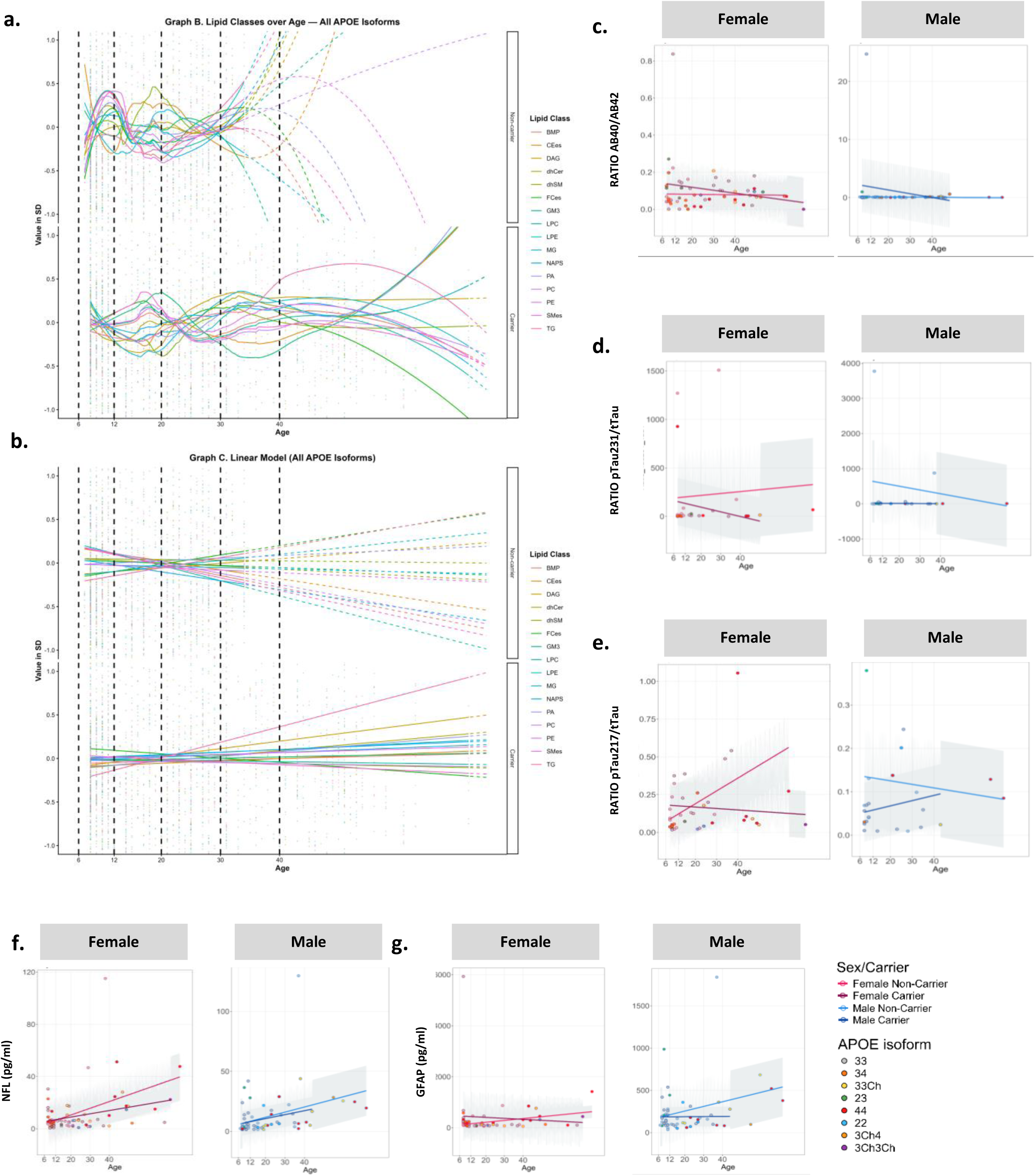
Age-dependent lipidome profiles and complementary biomarker ratios across APOE isoforms. **(a)** Smoothed LOESS regression curves (mean ± SD) showing age-dependent trajectories of the same 16 lipid classes in PSEN1-E280A non-carriers considering all APOE isoforms. **(b)** Linear model fits for lipid classes in PSEN1-E280A carriers (all APOE isoforms). **(c–g)** SIMOA serum biomarker trajectories (pg/m), shown separately for females and males, including **(c)** Aβ42/Aβ40, **(d)** pTau231/tTau ratio, **(e)** pTau217/tTau ratio, **(f)** NfL, and **(g)** GFAP. All SIMOA biomarkers (Aβ40, Aβ42, NfL, GFAP, pTau231, pTau217, and total tau) were analyzed across age using all APOE isoforms represented in the lipidomics subset.

**Extended Data Fig. 3.**
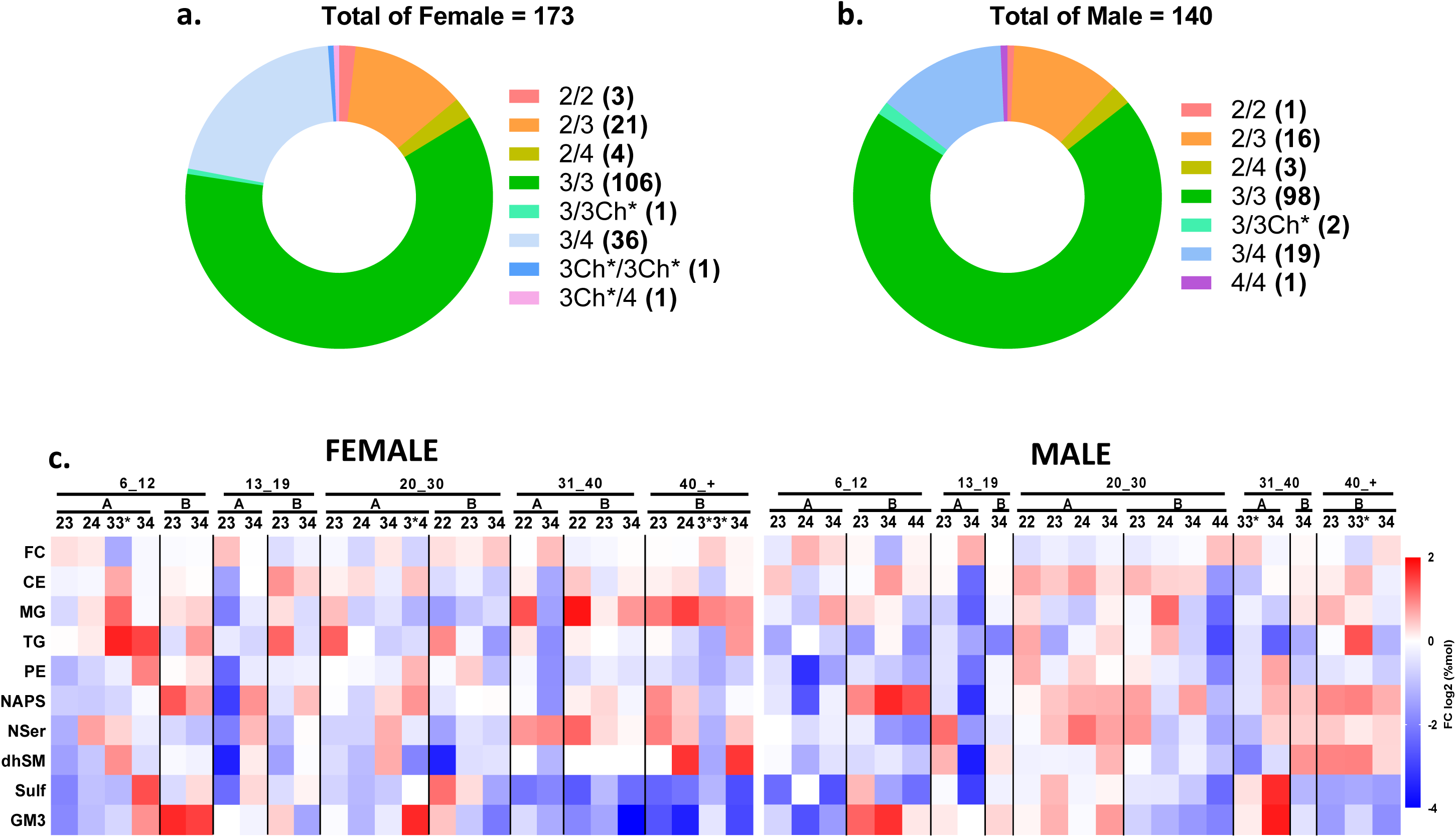
APOE isoform-dependent modulation of peripheral lipid trajectories. **(a)** Doughnut chart showing the distribution of APOE isoforms among female participants. **(b)** Doughnut chart showing the distribution of APOE isoforms among male participants. The same APOE isoforms were analyzed in both sexes: ε2/ε2, ε2/ε3, ε2/ε4, ε3/ε3, ε3/ε3*, ε3/ε4, ε3*/ε3*, ε3*/ε4, and ε4/ε4. APOE 3* indicates variants previously labeled as ‘Ch’ (Christ Church). **(c)** Heatmap showing age-dependent trajectories of lipid classes across APOE genotypes, stratified by sex and referenced to age– and sex-matched ε3/ε3 controls. Color scale indicates log₂ fold-change relative to controls (blue, negative; white, no change; red, positive). Panels A and B correspond to non-carriers and carriers, respectively, for each APOE isoform included. The lipid classes shown were selected based on the trajectory analyses performed exclusively in APOE3/3 individuals, which served as the reference framework for identifying age-sensitive metabolic patterns.

**Extended Data Fig. 4.**
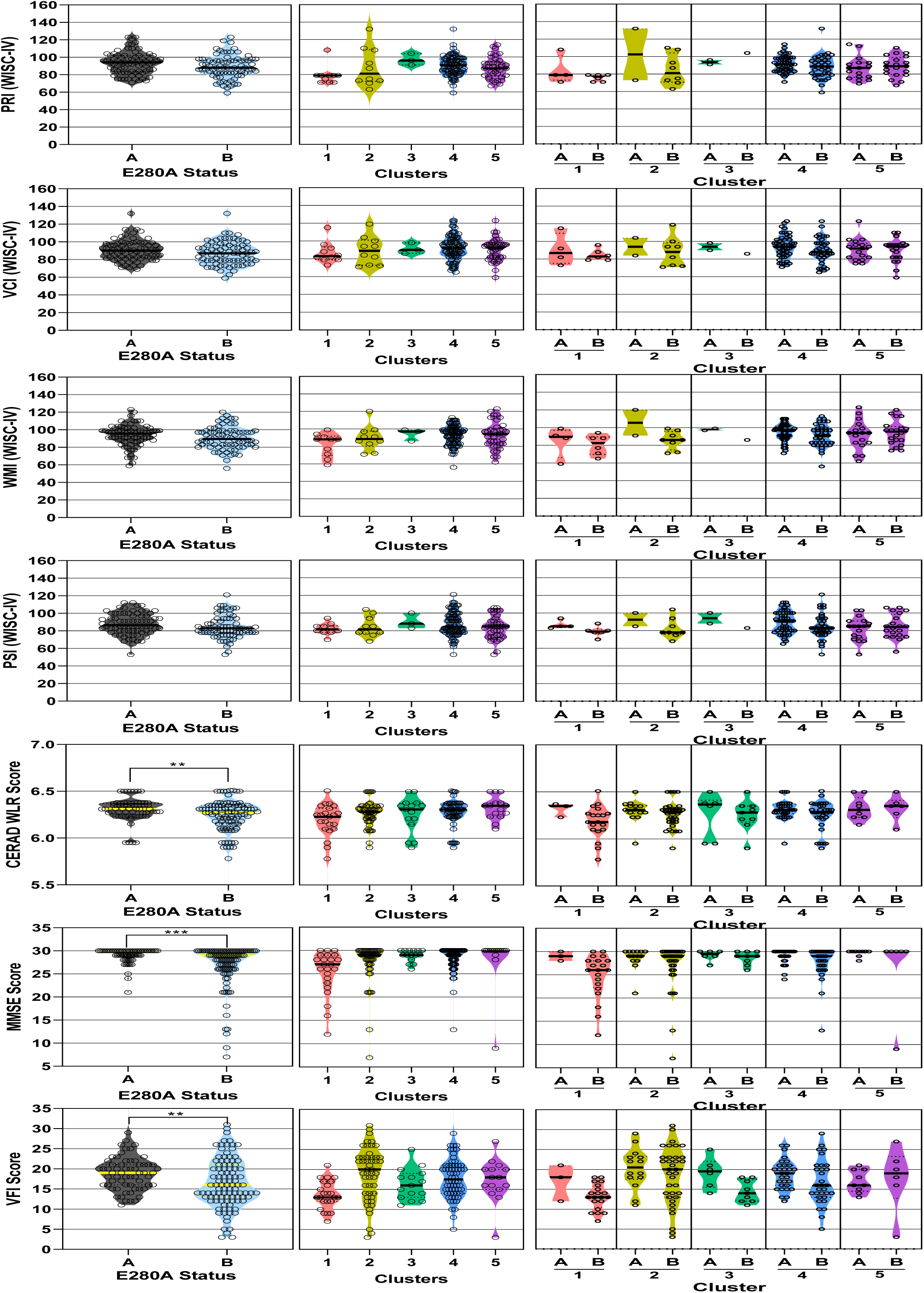
Neuropsychological performance across childhood, adolescence, and adulthood. This panel summarizes cognitive performance across age groups using standardized neuropsychological assessments. For children and adolescents, WISC-IV composite indices are shown, including: Verbal Comprehension Index (VCI), Perceptual Reasoning Index (PRI), Working Memory Index (WMI), and Processing Speed Index (PSI). For adults, cognitive performance is represented using the CERAD Word List Recall Score, Mini-Mental State Examination (MMSE) Score, and the Semantic Verbal FluencyScores. The first column shows violin plots displaying performance according to PSEN1-E280A carrier status (A: non-carriers; B: carriers), while the second and third columns show the distribution of cognitive scores across the LPA-derived clusters. cluster 1 (pink), cluster 2 (yellow), cluster 3 (green), cluster 4 (blue), cluster 5 (purple). Data is expressed in mean Z-Score according to age and years of education.

**Extended Data Fig. 5.**
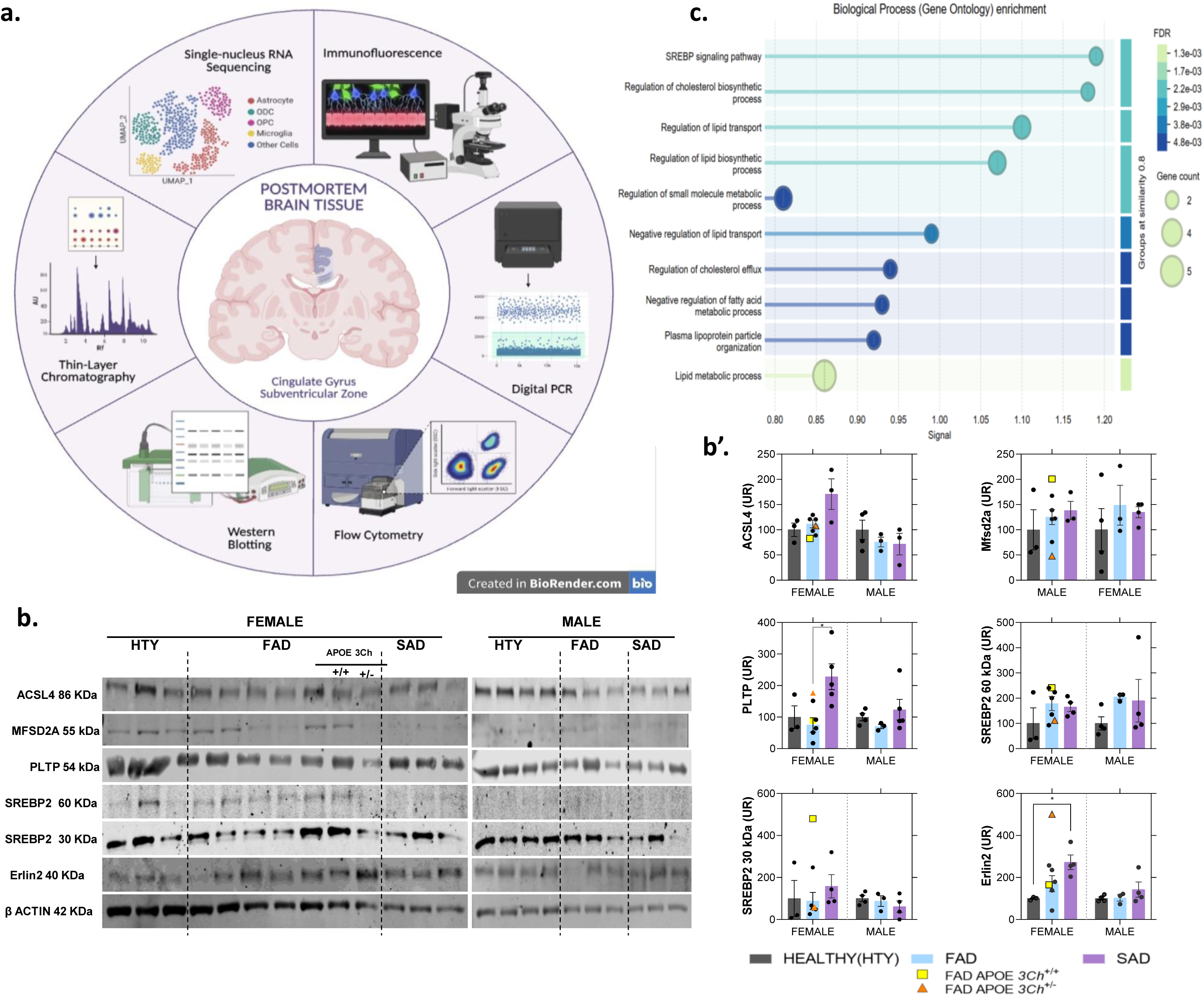
Postmortem sampling and lipid metabolism–related protein expression. **(a)** Schematic representation of postmortem tissue sampling (cingulate, SVZ and corpus callosum) and analytical approaches including immunofluorescence, dPCR, flow cytometry, western blotting, thin layer chromatography, snRNA-seq (created with BioRender.com) (**b) W**estern blots and **(b′)** corresponding quantifications of proteins involved in lipid regulation (Erlin2, SREBP2), transport (MFSD2A, PLTP), and fatty acid activation (ACSL4). Data are expressed as mean ± SEM; significance: *p < 0.05, **p < 0.01, ***p < 0.001 (N = 23). For women, groups include healthy controls (HTY, n = 3), FAD–PSEN1 E280A carriers (n = 5), FAD APOE3Ch+/+ carriers (n = 1), FAD APOE3Ch+/– heterozygous carriers (n = 1), and SAD cases (SAD, n = 3). For men, groups include HTY (n = 4), FAD–PSEN1 E280A carriers (n = 3), and SAD (n = 3). Parametric comparisons were performed using one-way ANOVA, whereas non-parametric data were analyzed using the Kruskal–Wallis test. **(c)** Biological process GO enrichment analysis based on the proteins analyzed, showing FDR values and gene counts.

**Extended Data Fig. 6.**
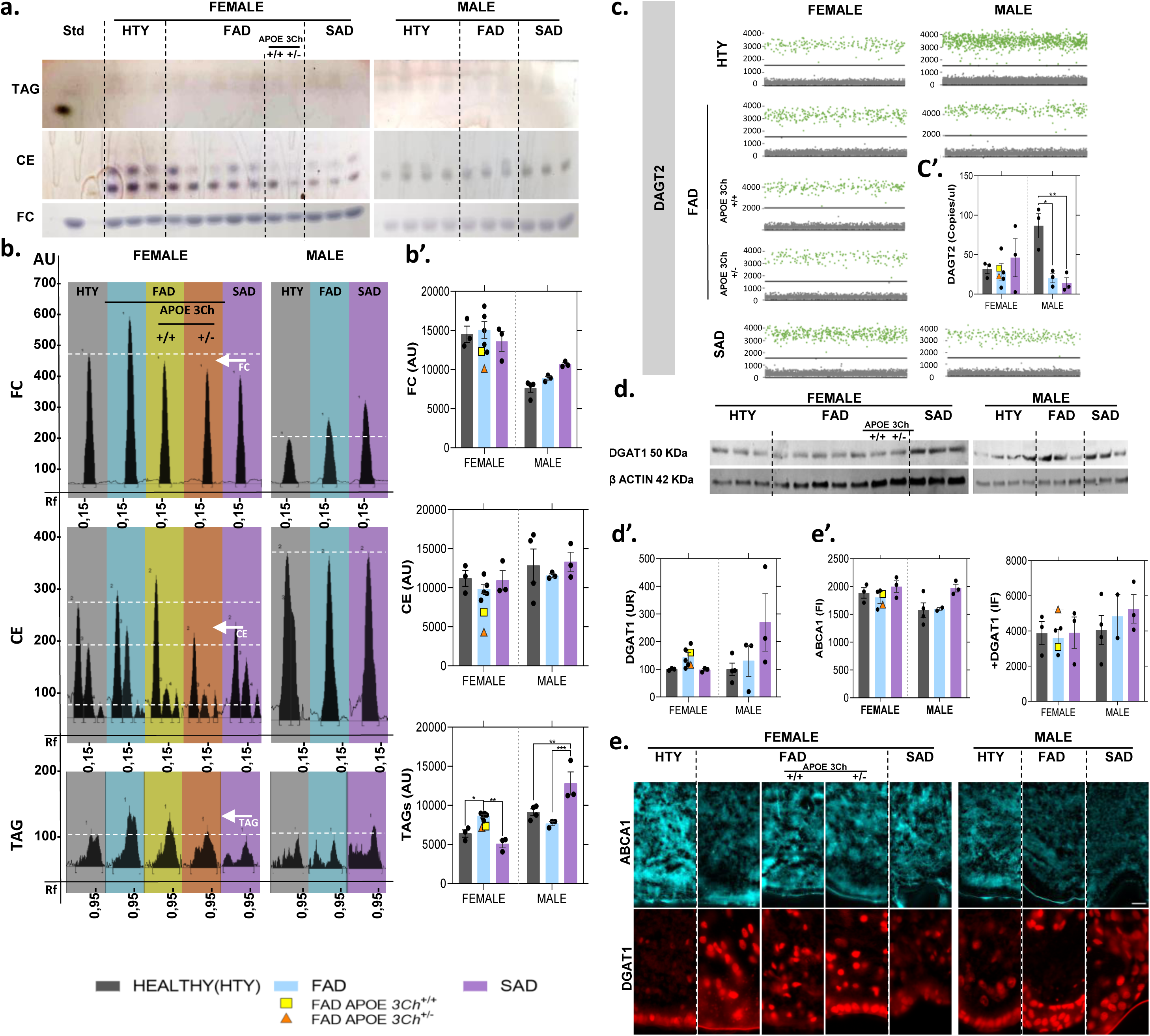
Analysis of neutral lipid metabolism in postmortem cingulate. **(a)** Representative thin-layer chromatography (TLC) plate showing the separation of neutral lipid classes, including free cholesterol (FC), cholesteryl esters (CE), and triacylglycerols (TAG). **(b–b′)** Representative TLC scan and corresponding quantifications of neutral lipid classes. **(c)** Representative scatter plots of individual dPCR reactions acquired using the ThermoFisher Q Studio dPCR platform, showing detection of DGAT2 mRNA in post-mortem cingulate tissue from women (left panel) and men (right panel). Each green dot corresponds to a positive droplet, defined by fluorescence above the automated threshold, whereas grey dots represent negative droplets. Individuals are grouped as healthy controls (HTY), carriers of FAD–PSEN1 E280A, carriers of FAD–PSEN1 E280A with the protective APOE3Ch+/+ variant, simple heterozygotes APOE3Ch+/-, and cases of sporadic Alzheimer’s disease (SAD). **(d) W**estern blots and **(d′)** Western blot quantification of DGAT1 protein levels. **(c’)** Bar plot showing the absolute quantification of DGAT2 (copies/µL) across diagnostic groups. **(e′)** Quantification of immunofluorescence (IF) signal in the SVZ of the cingulate cortex, showing localization patterns of DGAT1 and ABCA1. Data are expressed as mean ± SEM; significance: *p < 0.05, **p < 0.01, ***p < 0.001 (N = 23). For women, groups include healthy controls (HTY, n = 3), FAD–PSEN1 E280A carriers (n = 5), carriers with the protective FAD APOE3Ch+/+ variant (n = 1), FAD APOE3Ch+/– heterozygous carriers (n = 1), and sporadic AD cases (SAD, n = 3). For men, groups include HTY (n = 4), FAD–PSEN1 E280A (n = 3), and SAD (n = 3). Parametric comparisons were performed using one-way ANOVA, whereas non-parametric data were analyzed using the Kruskal–Wallis test.”

**Extended Data Fig. 7.**
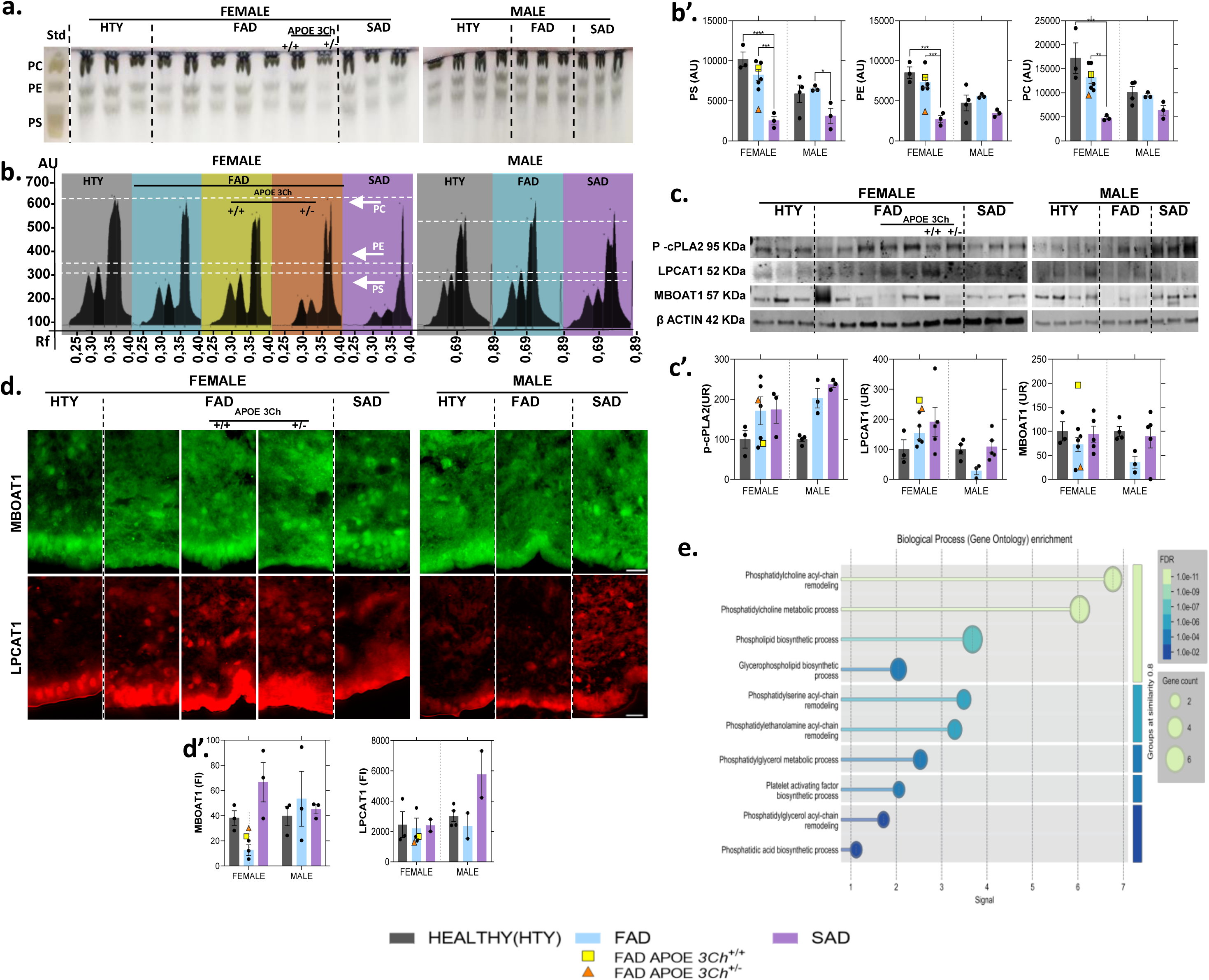
Analysis of phospholipid metabolism and remodeling enzymes in postmortem cingulate. **(a)** Representative thin-layer chromatography (TLC) plate showing the separation of major phospholipid classes, including phosphatidylserine (PS), phosphatidylethanolamine (PE), and phosphatidylcholine (PC). **(b–b′)** Representative TLC scan and corresponding quantification of phospholipid classes. **(c) W**estern blots and **(c′)** Western blot quantification of phospholipid remodeling enzymes, including cPLA₂, LPCAT1, and MBOAT1. **(d-d′)** immunofluorescence (IF) signal in cortical tissue, depicting MBOAT1 and LPCAT1 colocalization patterns and their Quantification. Data are expressed as mean ± SEM; significance: *p < 0.05, **p < 0.01, ***p < 0.001 (N = 23). For women, groups include healthy controls (HTY, n = 3), FAD–PSEN1 E280A carriers (n = 5), FAD APOE3Ch+/+ carriers (n = 1), FAD APOE3Ch+/– heterozygous carriers (n = 1), and sporadic AD cases (SAD, n = 3). For men, groups include HTY (n = 4), FAD–PSEN1 E280A carriers (n = 3), and SAD (n = 3). Parametric comparisons were performed using one-way ANOVA, whereas non-parametric data were analyzed using the Kruskal–Wallis test. **(e)** Gene Ontology (GO) enrichment analysis of biological processes associated with the analyzed proteins.

**Supplementary Figure 1. Pairwise correlation and distribution matrix of lipid classes**.

Comprehensive correlation matrix showing pairwise Pearson’s r values among all quantified lipid classes across the entire cohort. Upper panels display correlation coefficients, with positive correlations in red and negative in blue. Lower panels depict bivariate scatterplots with density distributions, highlighting co-regulation patterns among major lipid categories, including sterols, glycerides, phospholipids, sphingolipids, and glycosphingolipids. Histograms along the diagonal represent class-wise distributions. This structure served as the basis for defining co-regulated lipid modules used in subsequent latent and temporal modeling analyses.

**Supplementary Fig. 2 Distribution of SIMOA protein levels across the five LPA-derived clusters**.

Serum SIMOA biomarker concentrations (pg/mL) were normalized and expressed as percentages relative to the maximum observed value per analyte. Panels show the distribution of each biomarker across the five latent profile analysis (LPA)–derived clusters: **(a)** NfL, **(b)** GFAP, **(c)** Aβ40, **(d)** Aβ42, **(e)** total tau, **(f)** pTau231, and **(g)** pTau217.

**Supplementary Fig. 3 Cell-cycle profiles measured by PI signal in flow cytometry**.

**(a)** Representative histograms of propidium iodide (PI) fluorescence intensity showing DNA content distribution across cell-cycle phases. Phase assignment into G1, S, and G2/M was performed using the Dean–Jett–Fox cell-cycle modeling algorithm, which fits the DNA content histogram to estimate the proportion of cells in each phase. **(a′)** Quantification of cell-cycle phase distribution based on model-derived estimates. Data are expressed as mean ± SEM; significance: *p < 0.05, **p < 0.01, ***p < 0.001 (N=23). For women, groups include healthy controls (HTY, n= 2-3), FAD–PSEN1 E280A carriers (n=3-5), carriers with the protective APOE3Ch+/+ variant (n=1), APOE3Ch+/– heterozygous carriers (n=1), and sporadic AD cases (SAD, n=3). For men, groups include HTY (n = 4), FAD–PSEN1 E280A (n=3), and SAD (n=3). Parametric comparisons were performed using one-way ANOVA, whereas non-parametric data were analyzed using the Kruskal–Wallis test.

**Supplementary Table 1. Sociodemographic and Clinical Characteristics of Participants Stratified by APOE3 Genotype**

This table summarizes the sociodemographic, and clinical characteristics of participants whose blood samples were used for lipidomic analysis, grouped according to APOE3 genotype. The dataset includes information on sex distribution, chronological age, PSEN1-E280A carrier status, and clinical diagnosis related to Alzheimer’s disease. Educational attainment (years of formal education), presence of metabolic comorbidities, and body mass index (BMI). Categorical variables are presented as n (%), whereas continuous variables are reported as mean (SD).

**Supplementary Table 2. Sociodemographic Characteristics of Participants Included in the SIMOA Plasma Analysis**

This table summarizes the sociodemographic and genetic characteristics of the individuals whose plasma samples were analyzed using SIMOA assays. Reported variables include age at sampling, sex, and PSEN1-E280A carrier status, along with plasma concentrations of NfL, GFAP, Aβ40, Aβ42, total tau, p-Tau231, p-Tau217, and the corresponding derived ratios (p-Tau217/T-Tau, p-Tau231/T-Tau, Aβ42/Aβ40). Categorical variables are presented as n (%), and continuous variables are expressed as mean (SD). Group comparisons were conducted using the Kruskal–Walli’s rank-sum test for continuous variables and Fisher’s exact test for categorical variables. NA indicates data not available (N=173).

**Supplementary Table 3 Sociodemographic characteristics of the postmortem tissue sample**.

Groups are stratified by cognitive status, with APOECh carriers included within the FAD group. The table reports Age of Onset (AoO) and Age of Death (AoD) across groups, as well as National Institute of Aging neuropathological categorization and total brain weight. AoO and AoD are expressed in years, the postmortem interval in hours, and total brain weight in grams. Categorical variables are presented as n (%), whereas discrete and continuous variables are shown as mean (SD). NA indicates data not applicable.

**Supplementary Table 4 List of primary and secondary antibodies used in this study**.

This table summarizes all reagents and antibodies employed for immunofluorescence (IF) and Western blot (WB) experiments, SIMOA, including reagent type, source, catalog identifiers, and applications.

## ABBREVIATIONS AND DEFINITIONS

AC: Acyl-CoA
ACSL4: Acyl-CoA synthetase long-chain family member 4
Aß40: Amyloid-ß 1–40
Aß42: Amyloid-ß 1–42
ALDH3A2: Aldehyde dehydrogenase 3A2
APOE: Apolipoprotein E
APOE2: APOE e2 allele
APOE3/3: APOE e3/e3 genotype
APOE3Ch: APOE Christchurch variant
APOE4: APOE e4 allele
AoD: Age of death
AoO: Age of onset
AUC: Area under the curve
BMI: Body mass index
BMP: Bis(monoacylglycero)phosphate
CE: Cholesteryl ester
Cer: Ceramide
cPLA2: Cytosolic phospholipase A2
dPCR: Digital PCR
dhCer: Dihydroceramide
dhSM: Dihydrosphingomyelin
DG: Diacylglycerol
DGAT1: Diacylglycerol acyltransferase 1
DGAT2: Diacylglycerol acyltransferase 2
FAD: Familial Alzheimer’s disease
FC: Free cholesterol
GFAP: Glial fibrillary acidic protein
Gb3: Globoside Gb3
GM3: GM3 ganglioside
GO: Gene Ontology
HTY: Healthy young controls
IBA1: Ionized calcium-binding adapter molecule1
IF: Immunofluorescence
LacCer: Lactosylceramide
LPA: Latent profile analysis
LPC: Lysophosphatidylcholine
LPCAT1: Lysophosphatidylcholine acyltransferase 1
LPE: Lysophosphatidylethanolamine
MG: Monoacylglycerol
MBOAT1: Membrane-bound O-acyltransferase 1
MFSD2A: Major facilitator superfamily domain-containing protein 2A
MCI: Mild cognitive impairment
mhCer: Monohexosyl-ceramide
NAPE: N-acyl-phosphatidylethanolamine
NAPS: N-acyl-phosphatidylserine
NfL: Neurofilament light chain
NSer: N-acyl-serine
PCA: Principal component analysis
PC: Phosphatidylcholine
PCE: Plasmalogen PC
PE: Phosphatidylethanolamine
PEp: Plasmalogen PE
PG: Phosphatidylglycerol
PI: Propidium iodide / Phosphatidylinositol (context-dependent)
PLS-DA: Partial least squares discriminant analysis
PLTP: Phospholipid transfer protein
PMI: Postmortem interval
PRKD1: Protein kinase D1
PPM1L: Protein phosphatase 1L
PS: Phosphatidylserine
PSEN1: Presenilin-1
ROC: Receiver operating characteristic
SAD: Sporadic Alzheimer’s disease
SIMOA: Single molecule array
SL: Sphingolipids
SM: Sphingomyelin
SMPD2: Sphingomyelin phosphodiesterase 2
SMPD3: Sphingomyelin phosphodiesterase 3
snRNA-seq: Single-nucleus RNA sequencing
SREBP2: Sterol regulatory element-binding protein 2
SVZ: Subventricular zone
tTau: Total tau
TG: Triacylglycerol
TLC: Thin-layer chromatography
UGT8: UDP-galactose ceramide galactosyltransferase
VIM: Vimentin
WM: White matter

---

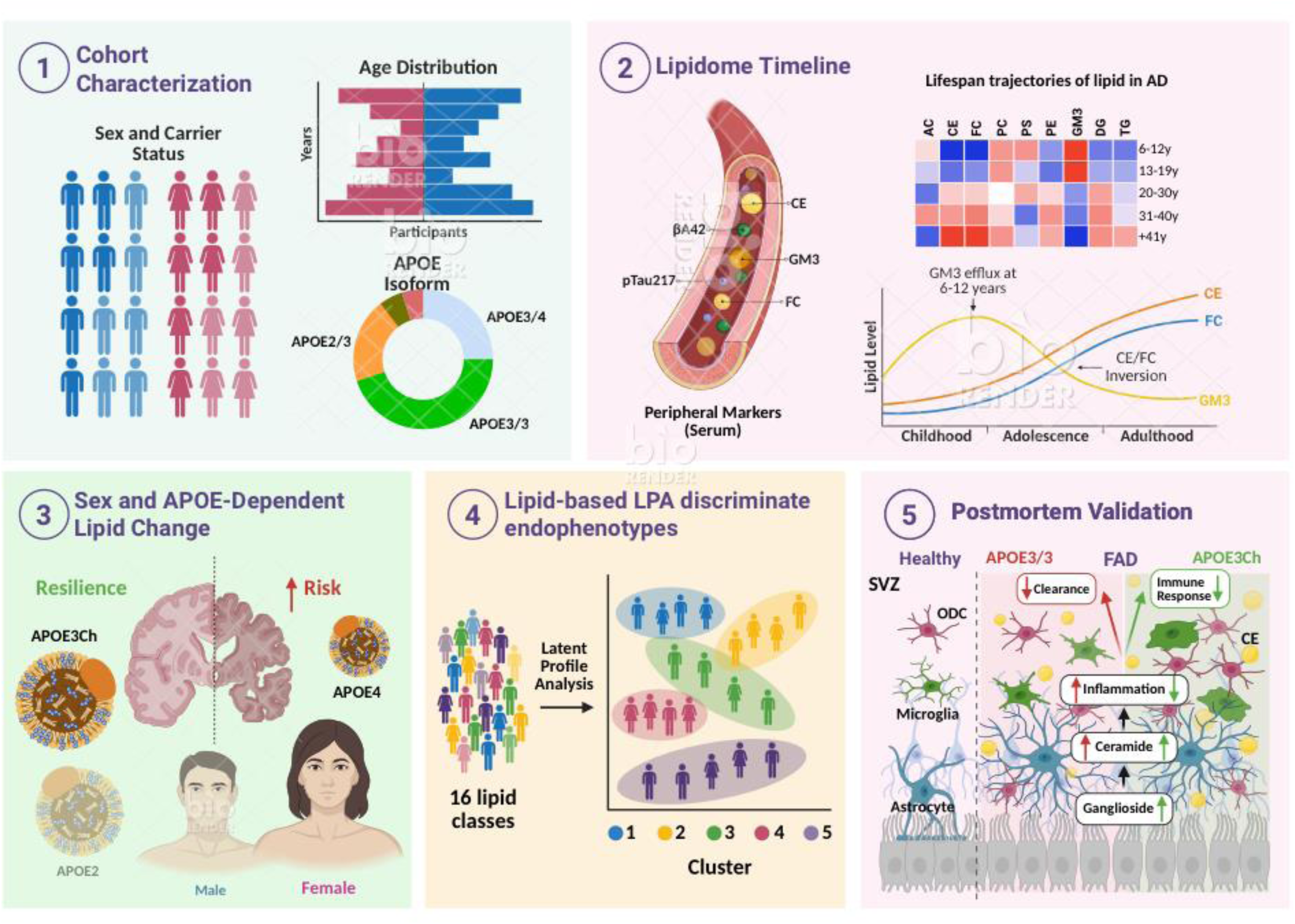

