## Supplementary material for "Serum Lipidome as an Early Peripheral Indicator in Familial Alzheimer’s Disease": Methods

**Study participants**

We retrospectively analyzed a cross-sectional dataset of 316 individuals selected from the historical Colombian PSEN1-E280A kindred registry. This registry, maintained by the Grupo de Neurociencias de Antioquia (GNA), encompasses over three decades of longitudinal follow-up for approximately 2,700 individuals. Participants were selected based on the availability of serum samples, comprehensive clinical assessments conducted between 2000 and 2020, and genetically confirmed PSEN1-E280A status. Not a priori sample-size calculation was performed. Three individuals were excluded due to confounding neurodegenerative or neurodevelopmental comorbidities.

To capture distinct neurodevelopmental and disease stages, participants were stratified into five age groups: childhood (6–12 years), adolescence (13–19 years), early adulthood (20–30 years), mid-adulthood (31–40 years), and late adulthood (≥41 years). APOE genotype was characterized, and PSEN1 carrier status was blinded to investigators (labeled as Groups A and B) during lipidomic analysis. The study adhered to ethical standards approved by the University of Antioquia’s Institutional Bioethics Review Committee, with written informed consent obtained from all participants or legal guardians (see Supplementary Table 1 for baseline characteristics).

**Clinical and sociodemographic variables**

Clinical and sociodemographic data were accrued via semi-structured interviews conducted by neurologists or general practitioners. Assessments encompassed socioeconomic status, educational attainment, medical history, and a comprehensive neurological examination. Diagnostic staging followed established criteria: Subjective Cognitive Decline (SCD) was defined by the presence of self-reported cognitive complaints in the absence of objective impairment on age- and education-adjusted neuropsychological assessments ^1^. Mild Cognitive Impairment (MCI) and dementia were diagnosed in accordance with DSM-5 guidelines ^2^. Functional independence was quantified using the Lawton–Brody Instrumental Activities of Daily Living (IADL) Scale ^3^ and the Barthel Index ^4^. Nutritional status was indexed by Body Mass Index (BMI), and comorbidities were verified through medical records and caregiver reports.

**Neuropsychological assessments**

Cognitive performance in adults was assessed using the Colombian validation of the Consortium to Establish a Registry for Alzheimer's Disease (CERAD) battery ^5^, covering memory, attention, language, praxis, and executive functions. Disease progression and functional severity were staged using the Lawton–Brody and Barthel indices, the Functional Assessment Staging Test (FAST) ^6^, and the Global Deterioration Scale (GDS) ^7^. Affective symptoms were screened using the Geriatric Depression Scale ^8^.

Pediatric participants (<18 years) followed the GNA’s standardized protocol, incorporating the Evaluación Neuropsicológica Infantil (ENI) ^9^ and the Behavior Assessment System for Children (BASC-2) ^10^ for behavioral profiling. General intellectual functioning was quantified via the Wechsler Intelligence Scale for Children (WISC-IV). In the absence of specific local norms at the time of assessment, scores were derived using the Chilean standardization ^11,12^, selected as the most demographically and linguistically proximal reference available for the Colombian population.

**Lipidomic analyses**

Serum samples (0.5 mL) from the Biobank of the Grupo de Neurociencias de Antioquia (GNA) were processed individually using the established Folch liquid-liquid extraction method ^13^. Lipids were extracted with a chloroform:methanol mixture (2:1 v/v) containing 0.005% butylated hydroxytoluene (BHT) as an antioxidant. Phase separation was achieved by homogenization, followed by the addition of 1 mL of 0.9% NaCl and centrifugation (3000 rpm, 3 min). The resulting organic phase was collected, evaporated to dryness under nitrogen, and lyophilized to ensure complete removal of residual moisture. Dried lipid extracts were subsequently subjected to mass spectrometric profiling. Quantitative lipidomic profiling was conducted via automated electrospray ionization tandem mass spectrometry (ESI–MS/MS) on an API 4000™/QTRAP 4000 platform (AB Sciex) at the Kansas Lipidomics Research Center (KLRC), following their standardized protocols ^14^. Each sample was spiked with internal standards corresponding to major lipid classes, including LPG(14:0), LPG(18:0), PG(14:0/14:0), LPE(14:0), LPE(18:0), LPC(13:0), LPC(19:0), PC(12:0/12:0), PC(24:1/24:1), LPA(14:0), LPA(18:0), PA(14:0), PA(20:0/20:0), PS(14:0/14:0), PS(20:0/20:0), PI(16:0/18:0), and PI(18:0/18:0). Lipid molecular species were identified based on their mass-to-charge ($m/z$) ratios and characteristic fragment ions. Quantification relative to internal standards was performed using the LipidomeDB Data Calculation Environment (<http://lipidome.bcf.ku.edu:8080/Lipidomics/index.jsp>). A total of 34 distinct lipid classes were quantitatively identified ^15^, with individual species defined by their total carbon number and degree of unsaturation. Lipid concentrations were normalized to molar percentages (%mol) across all detected species within each sample.

**Statistical analysis**

Statistical analyses were performed transversely within a retrospective design framework. Baseline characteristics were summarized as n (%) for categorical variables and mean ± SD for continuous variables. Group differences were assessed using Fisher’s exact test (categorical) and the Kruskal–Walli’s test (continuous), given that the Shapiro–Wilk test indicated a predominant non-Gaussian distribution across continuous variables. Lipid concentrations were calculated as the sum of total moles per class and normalized to molar percentage (%mol). For robust lipidomic analyses, only species detected in ≥80% of samples were included. To visualize lipidomic remodeling, heatmaps were generated using log2 fold-change (FC) values, referencing non-carriers harboring the APOE 3/3 genotype as the control group. For multiple comparisons, Kruskal–Wallis was followed by Dunn’s post hoc test, with p-values adjusted using the Benjamini–Hochberg False Discovery Rate (FDR) correction. Multivariate modeling captured global lipidomic variance. Principal Component Analysis (PCA) was employed as an unsupervised approach, and Partial Least Squares–Discriminant Analysis (PLS–DA) was implemented as a supervised dimensionality reduction method ^16^. Model quality was assessed via leave-one-out cross validation, with cluster separation defined by 95% confidence ellipsoids. Discriminant variables were identified using the Variable Importance in Projection (VIP) and Significance Multivariate Correlation (sMC) indices over the components provided by PLS-DA to minimize non-informative variance. Receiver Operating Characteristic (ROC) curves and Pearson’s correlation matrices were constructed to evaluate predictive accuracy and interclass dependencies, respectively.

Post-mortem brain tissue datasets were analyzed separately using GraphPad Prism v8.0. Comparisons across diagnostic categories (Healthy, sporadic AD, and familial AD) and sex-stratified analyses utilized one-way ANOVA and Tukey's post hoc tests, or appropriate non-parametric equivalents. Results are expressed as mean ± SEM, with statistical significance set at p < 0.05. Significance thresholds were defined as: p < 0.05 (*), $p < 0.01 (**), p < 0.001 (***), and p < 0.0001$ (****).

**Latent Profile Analysis**

Latent Profile Analysis (LPA) was employed as a model-based probabilistic clustering approach to identify unobserved (latent) subgroups within the lipidomic dataset. This method assumes that population heterogeneity can be represented by a finite mixture of continuous multivariate distributions, facilitating the detection of biologically meaningful endophenotypes not apparent through manifest variables alone. The LPA input matrix consisted exclusively of quantitative lipid species measurements, ensuring that clustering reflected intrinsic lipidomic variance rather than external covariates (genetic status, sociodemographic, or clinical scores were excluded from this analysis). Prior to model estimation, lipid classes included in the LPA were selected based on pairwise correlation analyses and inspection of distribution matrices, ensuring that only species with coherent covariance structures and non-redundant signal patterns contributed to latent class formation (Supp. Fig. 1). Models specifying one to nine latent classes were estimated using Gaussian finite mixture modeling assumptions in R (v4.4.1) within the RStudio environment (v2025.05.1), utilizing the Mclust package (version 6.1.1) ^17^.

Model selection was guided by the Bayesian Information Criterion (BIC), the Akaike Information Criterion (AIC), and entropy values, jointly optimizing model parsimony and the quality of the latent-profile clustering. The optimal number of profiles was determined as the model minimizing BIC while maintaining high entropy, which is indicative of stable cluster assignment probabilities. Beyond statistical fit indices, biological interpretability was incorporated as an essential selection criterion. This refinement prioritized profiles that captured coherent lipidomic trajectories and exhibited neurobiological relevance, ensuring class discrimination was meaningful. The resulting latent profiles delineate subgroups with internally homogeneous lipid distributions and distinct interclass separation patterns. This data-driven stratification provides a framework to evaluate whether lipidomic architecture can uncover early endophenotypes associated with Alzheimer’s disease (AD) risk, enabling the identification of metabolic substructures potentially reflecting early pathophysiological divergence for improved disease stratification and precision-targeted interventions.

**Simoa Biomarker Quantification and Quality Control**

Serum concentrations of neurofilament light chain (NfL), glial fibrillary acidic protein (GFAP), amyloid-β40 (Aβ40), amyloid-β42 (Aβ42), total tau (t-tau), and phosphorylated tau at threonine 231 (pTau231) and threonine 217 (pTau217) were measured using the SR-X Ultra-Sensitive Biomarker Detection System (Quanterix, Lexington, MA, USA) based on Single Molecule Array (Simoa) technology ([Simoa® Technology | Quanterix](https://www.quanterix.com/simoa-technology/)) ^18^. A total of N = 173 participants with available serum were included in this analysis (see Supplementary Table 2 for demographic characteristics and stratification by genotype, age group, and cognitive status). Analyses were performed according to manufacturer instructions using the following kits: Simoa Neurology 2-Plex B, Human Neurology 3-Plex A (N3PA), pTau-231 Advantage, and ALZpath pTau-217 Care Advantage.

The NfL–GFAP, Aβ40–Aβ42–t-tau, and pTau217 assays were conducted using a two-step protocol, whereas pTau231 followed a three-step protocol. Calibrators and serum samples were run in duplicate and diluted 1:4, except for pTau217 (1:3). Concentrations were derived from 4-parameter logistic (4-PL) calibration curves.

Each assay run includes low- and high-level quality controls (QC1 and QC2) to monitor inter-assay variability. Plate-specific correction factors were calculated from the ratio between the overall geometric mean of QC values and the observed value for each plate. Intra-assay and inter-assay coefficients of variation were maintained below 15% and 20%, respectively. When QC1 and QC2 deviated in opposite directions, a two-point anchoring correction based on the regression of expected versus observed values was applied. Measurements below the lower limit of quantification (LLOQ) were excluded from further analyses.

**Statistical Analysis for Simoa Data**

Statistical analyses were performed in R (v4.4.1) using RStudio (version 2025.05.1). Continuous variables were summarized as mean ± standard deviation or median (IQR), depending on distribution, and categorical variables as counts and percentages. Data cleaning and descriptive statistics were generated using the tidyverse suite, including dplyr and gtsummary ^19,20^.

Associations between serum biomarker concentrations and age were examined using linear models stratified by genetic status (PSEN1 mutation carriers vs. non-carriers) or demographic subgroups when appropriate. Group comparisons were conducted using the Mann–Whitney U test for two groups or the Kruskal–Wallis test for multiple age strata, followed by Dunn’s post hoc testing with Benjamini–Hochberg false discovery rate correction. Data visualizations were generated using ggplot2, and final layouts were assembled with the patchwork package.

**Postmortem Human Samples**

A total of 23 postmortem human brain samples were analyzed, including individuals with familial Alzheimer’s disease (FAD) carrying the PSEN1 E280A mutation, sporadic Alzheimer’s disease (SAD), and cognitively unimpaired controls. Two additional cases harboring the APOE-R154S (Christchurch) variant in one or both alleles were included for comparative analysis (Supplementary table 3). Neuropathological classification for all cases followed current Alzheimer’s disease diagnostic criteria ^21^.

Postmortem tissue, plasma, and serum were obtained with informed consent and repository authorization, in accordance with the Belmont Report and with approval from the Institutional Review Board of the Universidad de Antioquia. Blood samples were collected in EDTA tubes, processed immediately, and stored at −80 °C until analysis. Brains were obtained within 12 h of death. One hemisphere was snap-frozen at −80 °C for biochemical analyses. For this study, dissections targeted the cingulate cortex, including adjacent portions of the corpus callosum and the subventricular zone. Detailed demographic, genetic, and neuropathological characteristics are provided in Supplementary Table 3.

**Western Blotting**

Tissue samples comprising cingulate cortex together with the adjacent portion of the corpus callosum and the SVZ were homogenized in lysis buffer containing 150 mM NaCl, 20 mM Tris (pH 7.4), 10% glycerol, 1 mM EDTA, 1% NP-40 and protease inhibitors (1 mg/mL). Protein extracts (20 µg per lane) were denatured at 95 °C for 3 min in loading buffer (0.375 M Tris, pH 6.8; 50% glycerol; 10% SDS; 0.5 M DTT; 0.002% bromophenol blue) and separated by SDS–PAGE using the Mini-PROTEAN system (Bio-Rad) with low–molecular weight standards.

Proteins were transferred onto nitrocellulose (Bio-Rad 162-0115) or PVDF membranes (Bio-Rad 162-0177) using a Trans-Blot SD semi-dry transfer system (Bio-Rad) in Schafer–Nielsen buffer (48 mM Tris, 39 mM glycine, 20% methanol, pH 9.2) at 15 V for 50 min. Membranes were blocked in 5% nonfat dry milk prepared in TTBS (20 mM Tris–HCl, pH 7.5; 500 mM NaCl; 0.05% Tween-20) and incubated overnight at 4 °C with primary antibodies (listed in Supplementary Table 4).

Fluorescent secondary antibodies were used for detection, and imaging was performed with the Odyssey Infrared Imaging System (LI-COR Biosciences, software v5.2). Densitometric analyses were carried out using the same software, and protein abundance was normalized to actin and expressed relative to control samples.

**Immunofluorescence**

Free-floating tissue sections (30 µm) from the cingulate cortex, adjacent corpus callosum, and subventricular zone were obtained using a cryostat (Leica CM1850 UV). Sections were exposed to an LED light source for three consecutive nights to reduce autofluorescence, rinsed in 0.1 M phosphate buffer (PB), and subjected to antigen retrieval in citrate buffer (1×, pH 6.0; Master-Diagnostic MAD-004071R/D) at 95 °C for 20 min. After additional PB washes, tissues were incubated in 50 mM ammonium chloride for 15 min to quench residual aldehydes and permeabilized with 0.3% Triton X-100.

Sections were then washed and blocked for 1 h at room temperature in PB containing 0.3% Triton X-100 and 1% bovine serum albumin (BSA). Samples were incubated with primary antibodies for 72 h at 4 °C, washed, and incubated with the corresponding secondary antibodies for 2 h at room temperature. Antibody sources and dilutions are listed in Supplementary Table 4. After final washes, sections were mounted using fluorescence-compatible medium. Imaging was performed using an Olympus IX81 epifluorescence microscope. Quantitative analyses were performed in ImageJ (version 1.54).

**Lipid Extraction and Analysis by Thin-Layer Chromatography (TLC)**

All reagents, materials, and tissue homogenates were kept on ice or at 4 °C throughout the procedure to preserve lipid integrity. Brain tissue from the cingulate cortex, adjacent corpus callosum, and subventricular zone was homogenized in Vance buffer (225 mM mannitol, 25 mM HEPES–KOH, pH 7.4, 1 mM EGTA, 2 mM MgCl₂, 2 mM MnCl₂) at a 10:1 buffer-to-tissue ratio. When required, homogenates were centrifuged to remove insoluble debris, and supernatants were used for downstream processing. Protein concentrations were determined using Bradford or BCA assays and normalized to 0.5–2 mg/mL.

Total lipids were extracted using a modified Folch procedure. Briefly, 2 mL of chloroform:ethanol (2:1, v/v) were added to each sample and vortexed. Subsequently, 1 mL of 0.9% NaCl and 1 mL of chloroform:methanol (2:1, v/v) were added sequentially with additional vortexing. Samples were centrifuged (3,500 rpm, 7 min, 4 °C), and the lower organic phase was collected, dried under nitrogen or vacuum, and resuspended in chloroform to 15 mg/mL.

For ganglioside-enriched fractions, the aqueous phase from the Folch extraction was subjected to two additional extractions with butanol to recover glycolipids, followed by ethanol washes to remove residual salts ^22^. Combined butanolic extracts were dried and resuspended in chloroform:methanol (2:1, v/v).

For chromatographic separation, silica gel 60 aluminum-backed TLC plates (Merck) were prewashed with hexane and equilibrated in the appropriate mobile phase. Samples (15–20 µL at 15 mg/mL) were applied to each plate. Neutral lipids were resolved in hexane:diethyl ether:acetic acid (80:20:0.1, v/v/v); phospholipids in chloroform:methanol:acetic acid:water (50:30:8:4, v/v/v/v); sphingolipids in chloroform:methanol:0.25% KCl (60:30:8, v/v/v); and gangliosides in butanol:acetic acid:water (15.3:3.75:3.75, v/v/v). Plates were air-dried and visualized with anisaldehyde spray reagent. Chromatograms were digitized using a CAMAG TLC Scanner 3 ([TLC Visualizer 3 | CAMAG](https://camag.com/product/camag-tlc-visualizer-3/)), and absorbance was measured at 480 nm for semi-quantitative densitometric analysis, densitometric analysis was performed using visionCATS software (CAMAG).

**Sphingomyelinase Activity Assay and Thin-Layer Chromatography**

All solutions, materials, and tissue extracts were maintained on ice or at 4 °C and processed under minimal light exposure to preserve fluorescent products. Tissue from the cingulate cortex, adjacent corpus callosum, and subventricular zone was homogenized in Vance buffer (225 mM mannitol, 25 mM HEPES–KOH, pH 7.4, 1 mM EGTA, 2 mM MgCl₂, 2 mM MnCl₂) at a 10:1 buffer-to-tissue ratio. When required, homogenates were centrifuged to remove debris, and supernatants were collected. Protein concentration was determined by Bradford or BCA assays and adjusted to 0.5–2 mg/mL.

For sphingomyelinase activity, 100 µL of protein extract (500 µg of protein) were incubated with 200 µL of reaction buffer (Vance buffer supplemented with 2 mM MgCl₂, 2 mM MnCl₂, and 1 µM BODIPY–sphingomyelin. Reactions were carried out at 37 °C for 2 h in the dark. Lipids were extracted using a modified Folch procedure: 2 mL chloroform:ethanol (2:1, v/v) were added and vortexed, followed by the sequential addition of 1 mL 0.9% NaCl and 1 mL chloroform:methanol (2:1, v/v). Samples were centrifuged (3,500 rpm, 7 min, 4 °C), and the organic phase was collected, dried under nitrogen or vacuum, and resuspended in chloroform to 15 mg/mL.

TLC analyses were performed on silica gel 60 aluminum-backed plates (Merck), prewashed with hexane and equilibrated in the mobile phase. Samples were spotted alongside BODIPY–sphingomyelin and BODIPY–ceramide standards. Chromatographic separation was performed using chloroform:methanol:0.22% CaCl₂ (60:35:8, v/v/v). After development, fluorescent lipids were visualized using a CAMAG TLC Scanner 3 ([TLC Visualizer 3 | CAMAG](https://camag.com/product/camag-tlc-visualizer-3/)), and BODIPY emission was recorded at 510 nm. Densitometric analysis was performed using visionCATS software (CAMAG). Sphingomyelinase activity was calculated as the ratio of BODIPY–ceramide product to remaining BODIPY–sphingomyelin substrate.

**Tissue Dissociation and Flow Cytometry Analysis**

Cingulate cortex samples previously fixed in formaldehyde were washed three times with phosphate-buffered saline (PBS) for 5 min to remove residual fixative and then cut into small fragments (~1–2 mm³). Enzymatic dissociation was performed by incubating tissue fragments in PBS containing 0.1% collagenase type II (Sigma-Aldrich, C6885) and 0.01% DNase I (Sigma-Aldrich, DN25) at 37 °C for 60 min, with gentle agitation every 10 min. Mechanical dissociation was achieved by gentle pipetting using a sterile 200 µL tip. The resulting suspension was passed through a 40–70 µm cell strainer (Falcon/Corning) to eliminate debris. Cells were centrifuged at 400 × g for 10 min, the pellet was resuspended in PBS, and this washing step was repeated twice to ensure removal of residual enzymes. Total cell number was quantified using a Neubauer hemocytometer before staining.

For flow cytometry, cells were fixed and permeabilized in 0.1% Triton X-100 (Sigma-Aldrich, T8787) in PBS. Samples were incubated for 60 min at room temperature with the following primary antibodies (1:100 in PBS): Iba1 (Invitrogen, MA5-27726), GFAP (eBioscience, 14-9892-82), Ki-67 (Abcam, ab16667), and vimentin (Invitrogen, MA5-11883). After washing, cells were incubated with fluorophore-conjugated secondary antibodies (1:200 in PBS; Invitrogen) for 30 min at room temperature. Cell cycle assessment was performed by staining with propidium iodide (10–20 µg/mL; Sigma-Aldrich, P4170) for 15 min at room temperature. Samples were finally resuspended in PBS and analyzed on a BD LSR Fortessa flow cytometer (BD Biosciences), using identical acquisition settings for all samples. Human astrocytes and peripheral-blood macrophages fixed in 4% paraformaldehyde served as positive controls. Data analysis was conducted based on fluorescence intensity profiles to distinguish individual cellular populations.

**RNA Extraction and Digital PCR (dPCR) Quantification of DGAT2 and SMPD3 Expression**

Absolute gene expression quantification was performed using digital PCR (dPCR) on the QuantStudio™ Absolute Q™ Digital PCR System (Thermo Fisher Scientific, USA). Total RNA was extracted from human cingulate cortex samples using the MagMAX™ mirVana™ Total RNA Isolation Kit (Thermo Fisher Scientific, A27828) following the manufacturer’s instructions. RNA purity and integrity were assessed spectrophotometrically, and concentrations were adjusted to 3.5 ng/µL for downstream dPCR reactions.

One-step reverse transcription and amplification were performed using the Absolute Q™ 1-Step RT-dPCR Master Mix (4X) (Thermo Fisher Scientific, A55146). TaqMan® Gene Expression Assays were used for DGAT2 (Assay ID Hs01045907_m1, VIC-labeled) and SMPD3 (Assay ID Hs00920354_m1, ABY-labeled). Reactions were loaded into Absolute Q™ Digital PCR chips and processed according to the manufacturer’s workflow for partition generation, thermal cycling, and fluorescence acquisition.

Data were analyzed using the Absolute Q™ Analysis Suite v6.3.4, applying automated thresholding for partition-based quantification. Gene expression values were reported as absolute copy number (copies/µL). All samples were run in technical duplicates, and non-template controls were included to exclude contamination or nonspecific amplification.

**Single-nucleus RNA Sequencing (snRNA-seq) Analysis**

Publicly available single-nucleus RNA sequencing (snRNA-seq) datasets from sporadic Alzheimer’s disease (SAD), familial Alzheimer’s disease (FAD), and APOE genotype-controlled cohorts were analyzed (GEO accessions: GSE222494, GSE222495, GSE206744). Raw data were processed using CellBender v0.2 (PMID: 37592183) to remove ambient RNA and identify high-quality nuclei via the remove-background tool. Downstream preprocessing, quality control, and normalization steps were performed using Scanpy v1.6.0 and AnnData v0.7.4 (PMID: 29409532).

Nuclei and genes were filtered according to the following criteria: (i) doublet exclusion with Scrublet v0.2.2 (10 principal components); (ii) nuclei expressing <300 or >7,000 genes; (iii) genes detected in fewer than 100 nuclei across all batches; (iv) genes with <5 total counts; (v) nuclei with >5% mitochondrial gene content; and (vi) removal of mitochondrial genes and the highly expressed lncRNA MALAT1. Batch correction was performed using Harmony v0.1.0 (PMID: 31740819) with default parameters. Normalized counts were converted to Counts Per Million (CPM) and log-transformed (log(CPM + 1)). The 2,000 most variable genes were selected, and dimensionality reduction was achieved by principal component analysis (PCA), retaining components explaining ≥90% of the total variance. A neighborhood graph was constructed using pp.neighbors (100 neighbors, 27 PCs), followed by UMAP embedding (PMID: 30531897). Clustering was performed using the Leiden algorithm at a resolution of 0.15 (PMID: 30914743). Cell-type annotation was conducted by integrating canonical marker genes with curated databases including CellMarker 2.0, Tabula Sapiens, PanglaoDB, and Azimuth. Subclustering analyses focused on astrocytes, microglia, oligodendrocytes, and oligodendrocyte precursor cells (OPCs). Differential gene expression (DGE) analyses were conducted using a pseudo-bulk approach implemented with EdgeR-LTR, comparing expression profiles between groups. Fold changes were computed relative to the mean expression of the reference group. Genes involved in lipid metabolism were prioritized, and differential expression was defined by p ≤ 0.05. Finally, to control for cell cycle heterogeneity, phase-specific scores (G1, S, and G2/M) were computed using Scanpy’s tl.score_genes_cell_cycle function, based on canonical markers such as CDK1, CDK4, MCM6, and PCNA (PMID: 27124452). All analyses were performed in R v4.3.2 and Python v3.8

**Protein–protein interaction network analysis.**

Protein–protein associations were obtained from the STRING database (accessed 23 November 2025; (<https://string-db.org>). We imported the list of differentially represented proteins (gene names / UniProt IDs) and generated organism-specific networks including both physical and functional associations, using the “full STRING network” view. Interactions were filtered at a minimum combined confidence score ≥0.70 (high confidence) and displayed edge weights corresponding to STRING’s combined score (which integrates experimental data, curated databases, text mining and other evidence channels). Networks were exported in tabular form (edge list and node attributes) via STRING’s “Tables/Exports” function for downstream analysis and visualization. Clustering of network modules was performed with the Markov Cluster Algorithm (MCL) implemented in STRING/Cytoscape (inflation = 2.0) to identify highly interconnected protein groups. Functional enrichment (Gene Ontology biological process, KEGG pathways, Pfam/InterPro domains and UniProt keywords) was performed using STRING’s built-in over-representation test (hypergeometric test) and p-values were adjusted for multiple testing by the Benjamini–Hochberg procedure; terms with FDR ≤ 0.05 were considered significant. For programmatic reproducibility we confirmed results using the STRING API / STRINGdb package and archived the exact query outputs (edge lists, enrichment tables and parameters) as source data.
