## Supplementary figures and images for "Serum Lipidome as an Early Peripheral Indicator in Familial Alzheimer’s Disease"

### Suplementary figure 2

#### Slide 1
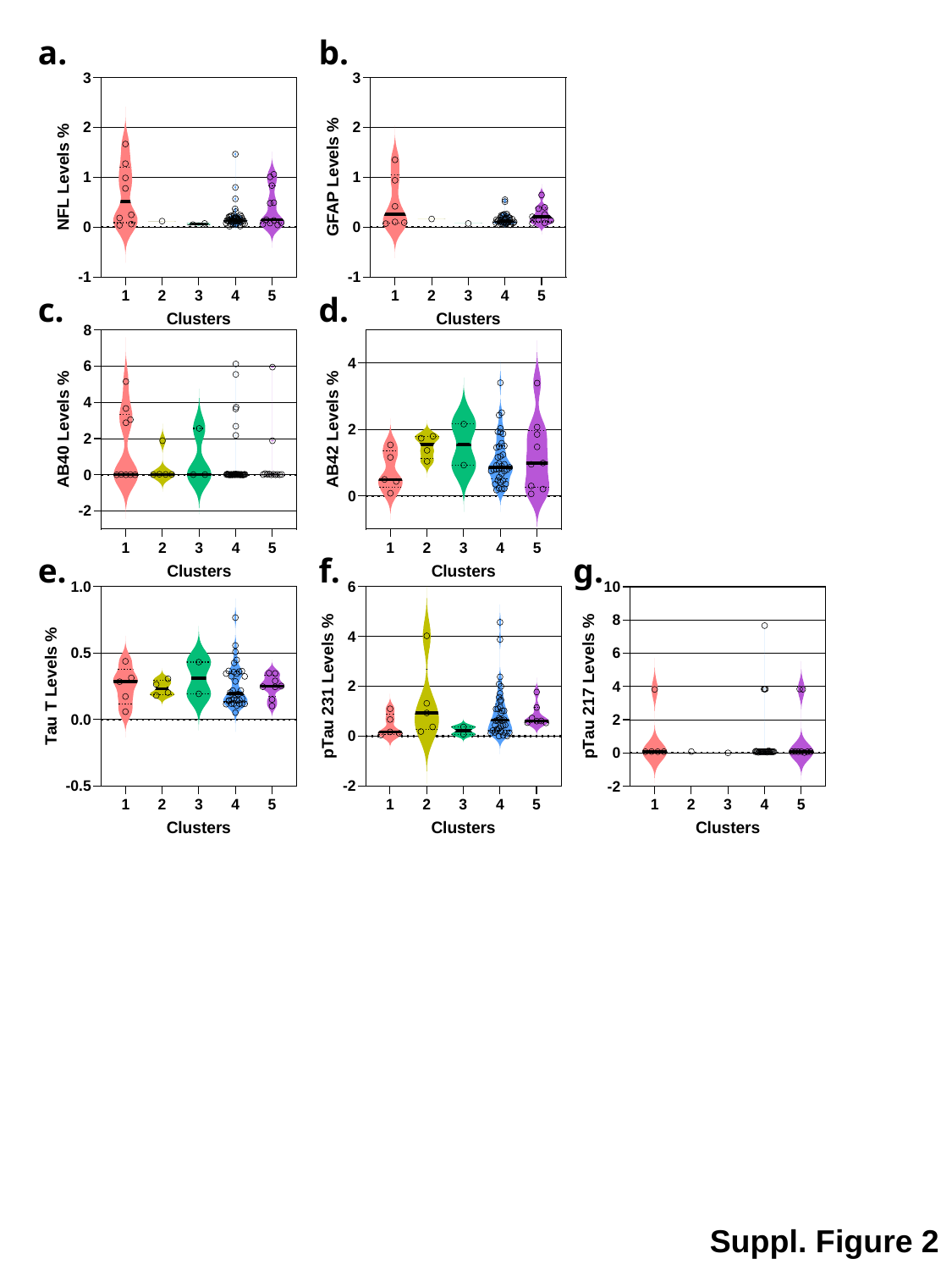

a.
b.
c.
d.
e.
f.
g.
Suppl. Figure 2

### Supplementary Figure 1

#### Slide 1
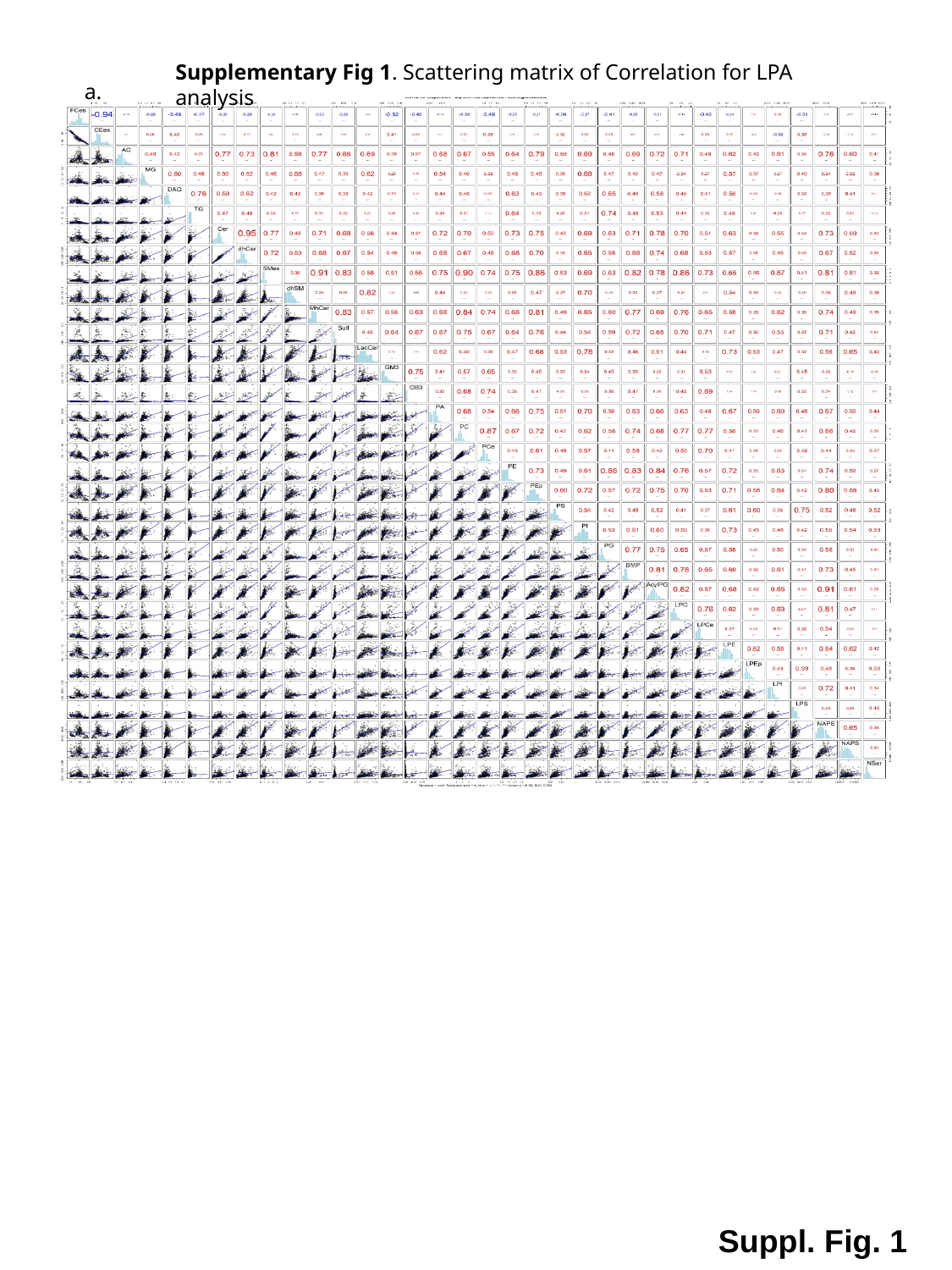

Supplementary Fig 1. Scattering matrix of Correlation for LPA analysis
a.
Suppl. Fig. 1

### Supplementary figure 3

#### Slide 1
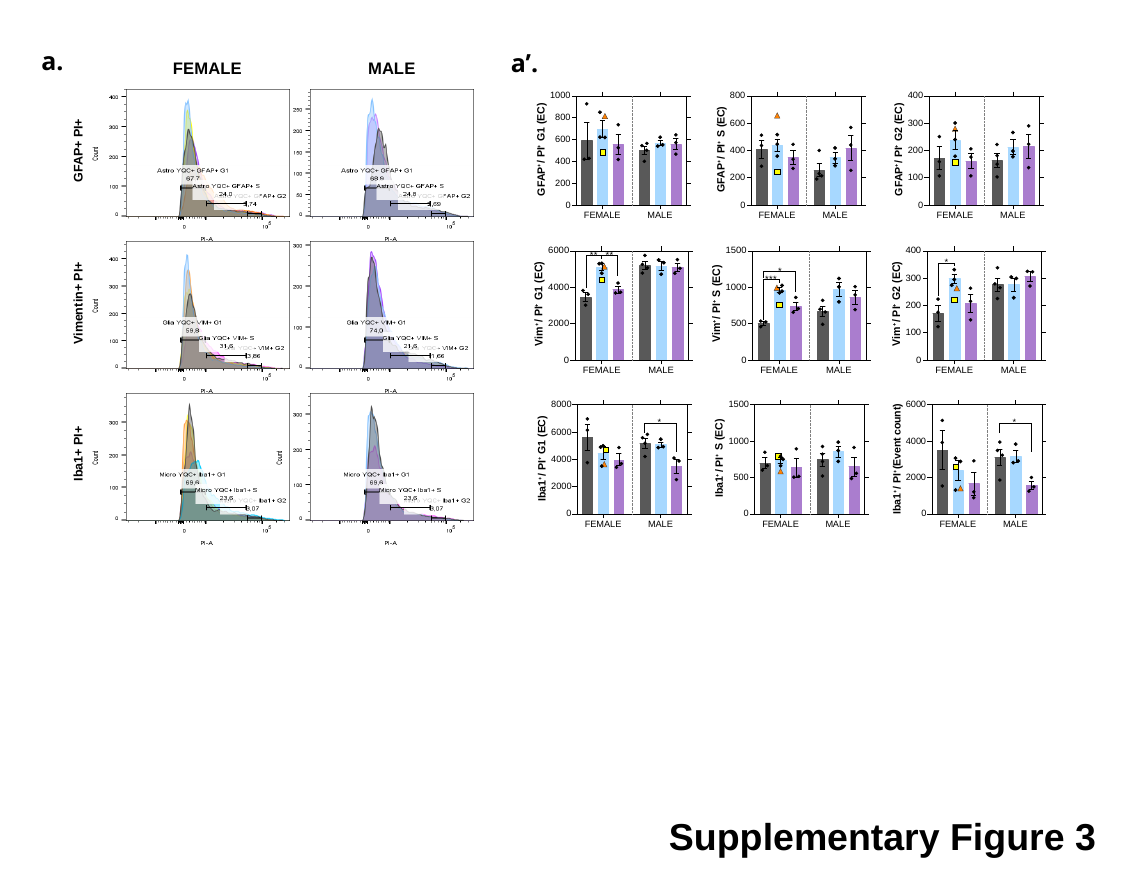

a.
a’.
FEMALE
MALE
 GFAP+ PI+
 Vimentin+ PI+
 Iba1+ PI+
Supplementary Figure 3

### Supplementary table 1

#### Slide 1
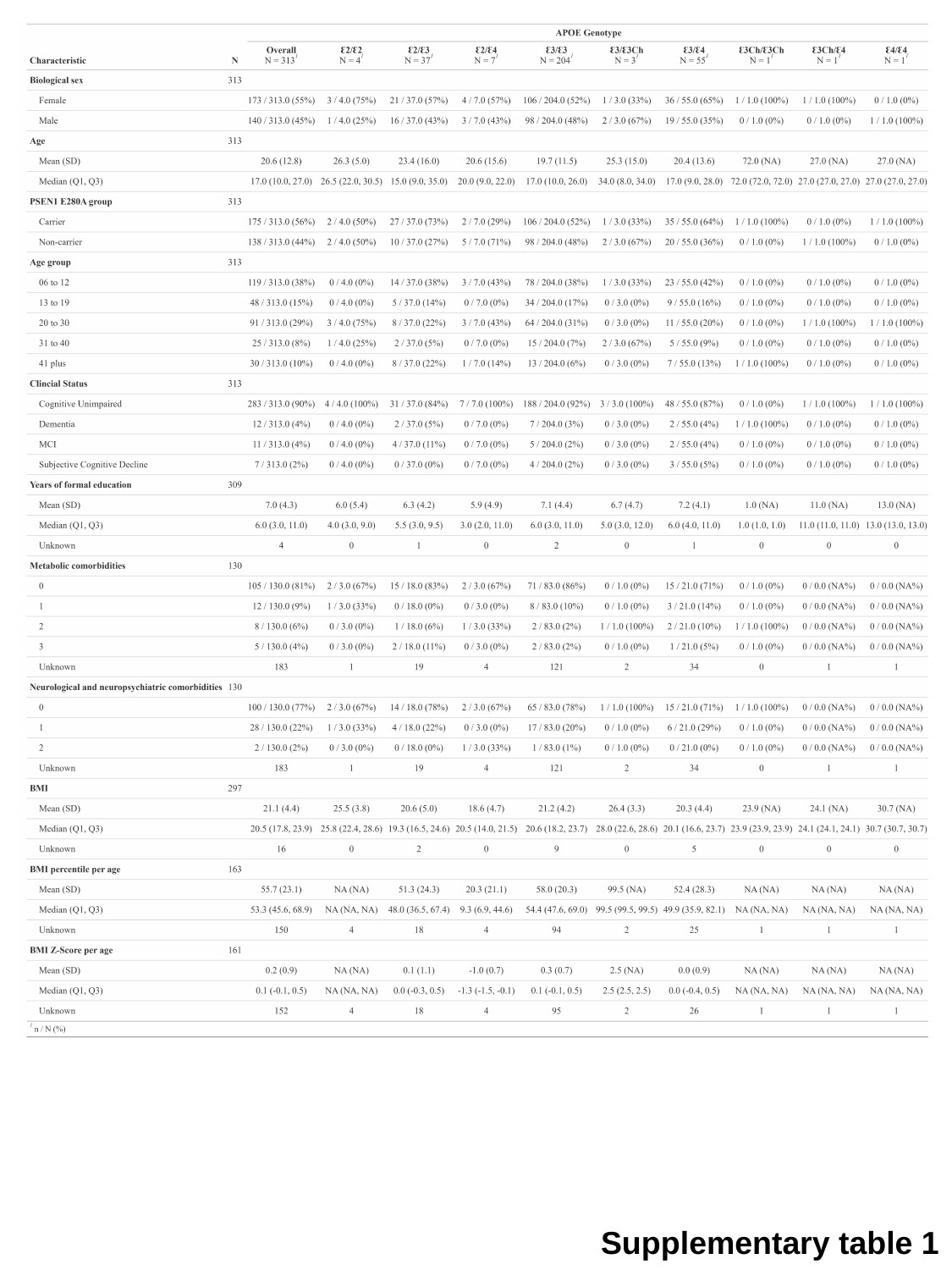

Supplementary table 1

### Supplementary table 2

#### Slide 1
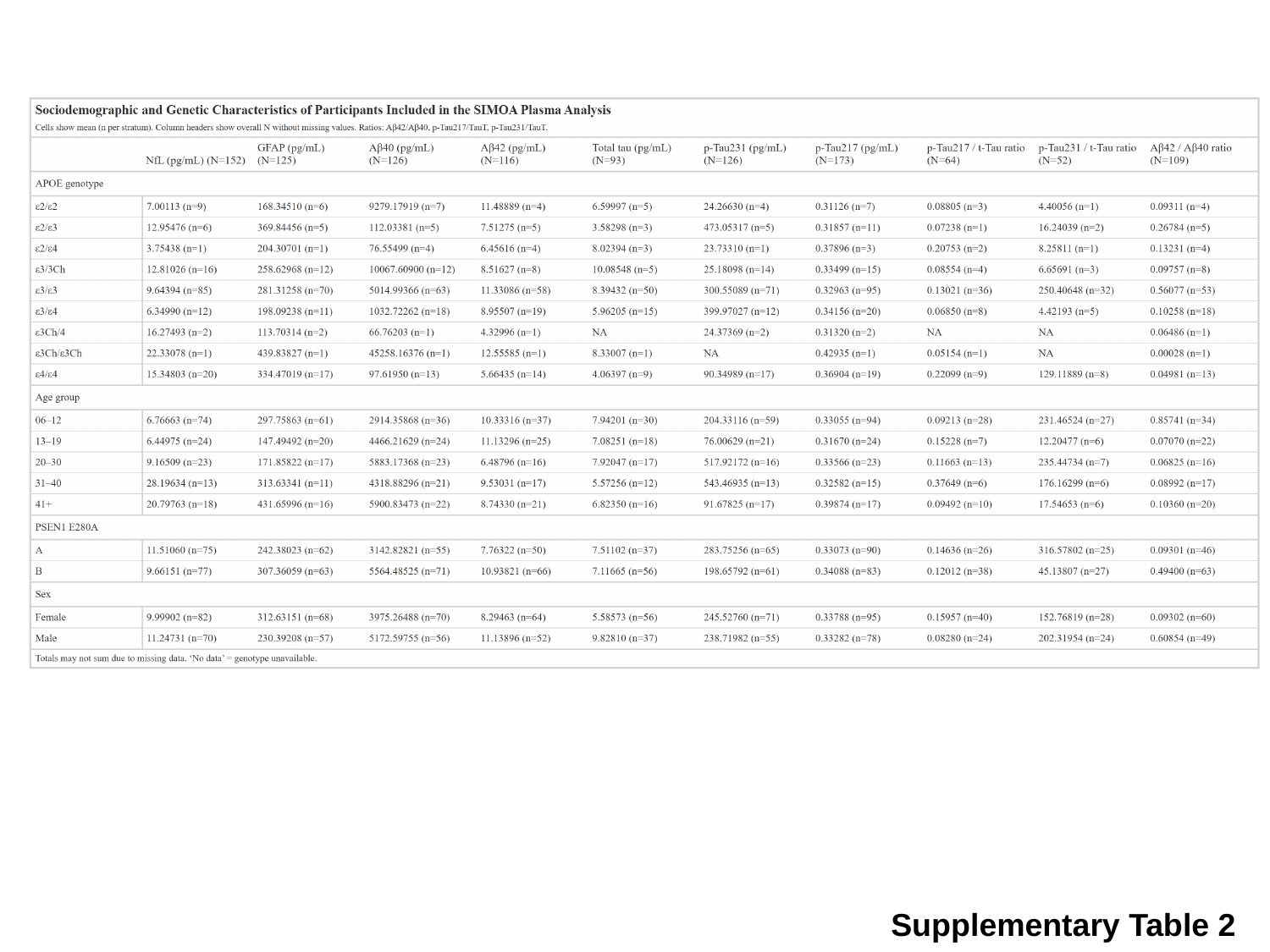

Supplementary Table 2

### Supplementary table 3

#### Slide 1
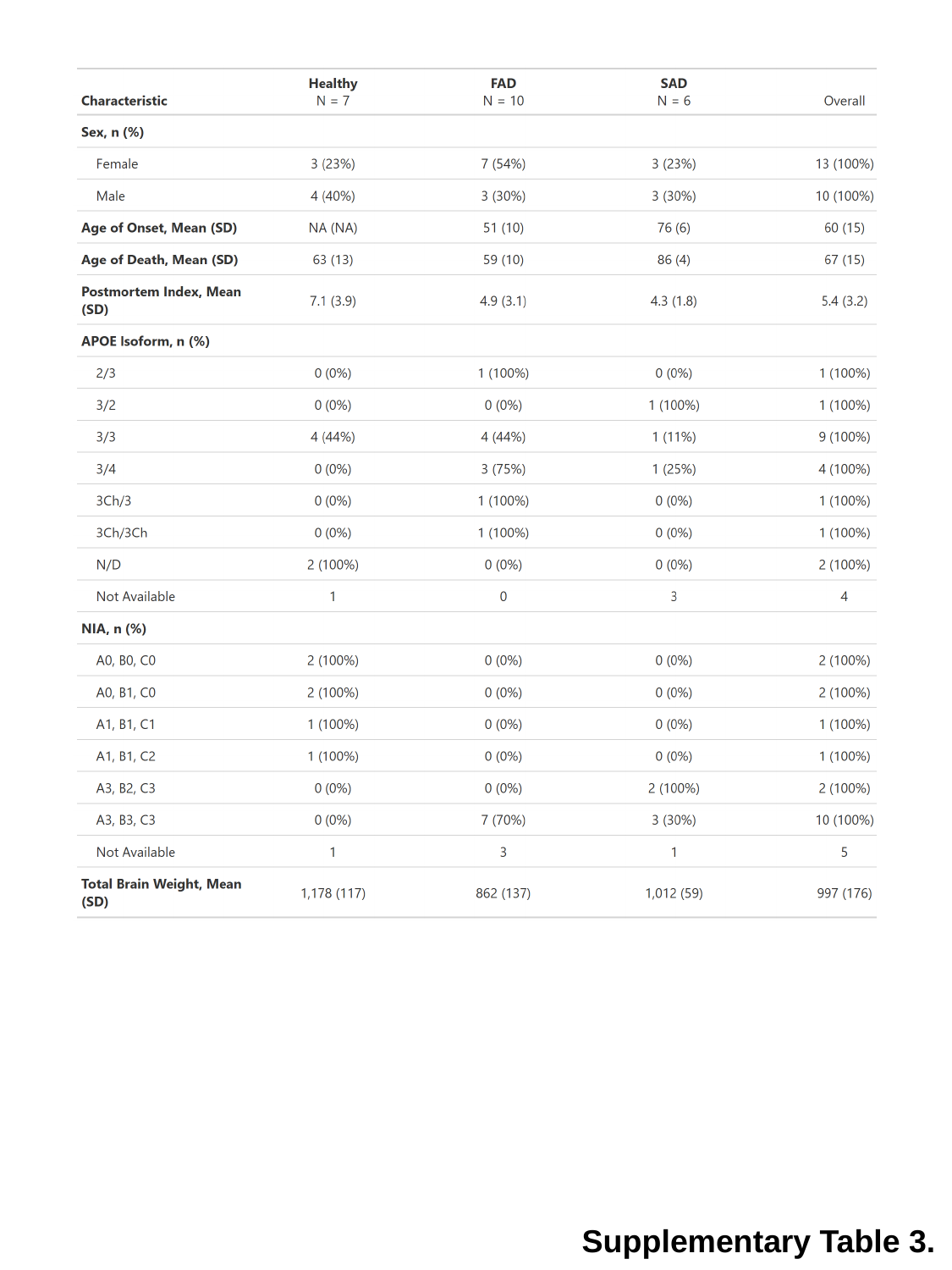

Supplementary Table 3.
